# Metabolic adaptation to maternal hyperglycemia via ACLY-dependent acetyl-CoA production drives epigenetic remodeling and dysregulated placental development

**DOI:** 10.64898/2026.03.17.712507

**Authors:** Mengying Liu, Keer Jin, Shihong Qi, Dandan Chen, Yichen Han, Weina Xu, Caihe Wen, Haiyan Wen, Yuanwei Liu, Bo He, Xiaona Lin

**Author notes:** Correspondence: M.L.,; B.H.,; X.L. M.L., K.J. and S.Q. contributed equally to this work.

## Abstract

Gestational diabetes mellitus (GDM) is a common metabolic complication of pregnancy that is paradoxically associated with both fetal overgrowth and fetal growth restriction (FGR). While maternal hyperglycemia is widely presumed to drive macrosomia through excessive nutrient supply, the mechanisms underlying FGR remain poorly understood. Here, using a mouse model that recapitulates the small-for-gestational-age (SGA) phenotype observed in human GDM pregnancies, we identify placental underdevelopment as a principal driver of FGR. Despite systemic nutrient abundance, hyperglycemic placentas exhibit reduced mass and an increased fetal-to-placental weight ratio, indicative of placental insufficiency. Mechanistically, maternal hyperglycemia induces anabolic metabolic rewiring while suppressing oxidative phosphorylation (OXPHOS), accompanied by upregulation and nuclear redistribution of ATP-citrate lyase (ACLY). ACLY converts glucose-derived carbon into acetyl-CoA in the cytosol and nucleus, thereby coupling glycolytic flux to lipid and hexosamine biosynthesis as well as to global histone hyperacetylation. This hyperacetylation-associated epigenetic reprogramming activates metabolic, innate immune, and inflammatory gene programs while repressing pro-proliferative and anti-apoptotic pathways. Consequently, placental growth is compromised despite nutrient excess. Importantly, activation of the ACLY-acetyl-CoA axis and global histone hyperacetylation is consistently observed in human GDM placentas across diverse birth outcomes, suggesting a conserved metabolic-epigenetic adaptation to maternal hyperglycemia. Together, these findings identify ACLY-dependent acetyl-CoA production as a central metabolic node linking maternal hyperglycemia to chromatin remodeling and placental development control, thereby reshaping fetal growth trajectories.

## Introduction

Gestational diabetes mellitus (GDM) is one of the most common medical complications of pregnancy, affecting approximately 14% of pregnant women worldwide (1). GDM is defined as impaired glucose tolerance that first arises or is first recognized during the second or third trimester (24-28 weeks) and is characterized by maternal hyperglycemia(2). Prolonged or uncontrolled hyperglycemia in GDM increases the risk of adverse birth outcomes, including large-for-gestational-age (LGA) infants, small-for-gestational-age (SGA) infants, and appropriate-for-gestational-age (AGA) infants with additional complications(3–5). Despite these associations, the mechanisms underlying the heterogeneity of birth outcomes in GDM remain largely unknown.

The placenta, a critical organ connecting the mother and fetus, mediates nutrient transport and gas exchange and regulates endocrine functions, thereby playing a pivotal role in supporting fetal development(6). Structural or functional abnormalities of the placenta can lead to adverse birth outcomes(6). Comparative studies of placentas from GDM patients and healthy controls have shown that GDM placentas are generally larger and heavier, likely reflecting increased nutrient availability under maternal hyperglycemia(7, 8). Accompanying this placental enlargement, GDM pregnancies are associated with a higher risk of LGA infants, while the fetal-to-placental weight ratio is significantly lower than that in non-GDM pregnancies(7). This disproportionate increase in placental and fetal mass suggests that each unit of placental tissue fails to adequately support the corresponding fetal growth, consistent with reports that maternal hyperglycemia may induce structural or functional placental insufficiency(9, 10). Indeed, studies in GDM mouse models have demonstrated that maternal hyperglycemia can cause both structural and functional placental abnormalities, particularly when hyperglycemia is severe or sustained(11–13). In addition to LGA infants, approximately 7% of GDM pregnancies result in fetal growth restriction (FGR) and SGA infants (3, 14). In Asian populations, the incidence of GDM-associated SGA is even higher, increasing from approximately 12.26% to 14.35% from 2014 to 2020(15). This paradoxical occurrence of SGA despite excessive nutrient availability remains poorly understood. Some studies suggest that severe placental insufficiency and increased placental inflammation may limit nutrient supply and contribute to FGR(5, 16), but mechanistic evidence supporting this hypothesis remains limited.

Emerging evidence suggests that appropriate glucose metabolism and acetyl-CoA production are essential for proper placental development, partly through the maintenance of histone acetylation(17, 18). In GDM, placental metabolic imbalance—particularly in lipid metabolism—has been widely reported(19, 20). Concurrently, the expression of metabolism-related and nutrient transporter genes is also altered, indicating that the GDM placenta undergoes extensive metabolic adaptation to hyperglycemia(11). Interestingly, several studies have also documented epigenetic alterations in GDM placentas, including changes in histone acetylation and DNA methylation(21–23). Given that abnormal lipid metabolism may stem from disrupted acetyl-CoA metabolism under hyperglycemia and considering that acetyl-CoA serves as a key substrate for histone acetylation(24, 25), the potential link between acetyl-CoA imbalance and epigenetic alterations in GDM placentas remains largely unclear. Furthermore, whether this metabolic-epigenetic rewiring contributes to placental dysfunction in GDM has yet to be elucidated.

Here, we established a mouse model that recapitulates the SGA phenotype observed in human GDM pregnancies. Using this model, we confirmed the well-established concept that maternal hyperglycemia markedly promotes anabolic metabolism, thereby increasing nutrient availability. Importantly, our findings further reveal that ACLY-dependent acetyl-CoA production represents a key mechanism by which the placenta metabolically adapts to maternal hyperglycemia in both mouse and human GDM. As a central metabolic intermediate, acetyl-CoA not only fuels lipid and hexosamine biosynthesis but also drives histone H3 hyperacetylation. This metabolite-driven epigenetic reprogramming reshapes gene expression programs that ultimately determine placental and fetus developmental fate.

## Results

### Gestational hyperglycemia induces FGR through impairing placenta development

To investigate how hyperglycemia affects placental development, we established a mouse model of GDM during late gestation, as previously described(13). Briefely, pregnant mice received low-dose streptozotocin (STZ) treatment on D6 (gestational day 1 refers to D1) and D12 to induce pancreatic β-cell dysfunction (***SI Appendix*, Fig. S1*A***). As expected, STZ-treated mice exhibited significantly elevated blood glucose levels during mid-to-late gestation compared with controls (***SI Appendix*, Fig. S1*B***). Glucose tolerance tests further confirmed impaired glucose handling in STZ-treated mice at late gestation (***SI Appendix*, Fig. S1*C***). Together, these results verify that STZ treatment successfully impairs pancreatic β-cell function and induces a GDM-like hyperglycemic state in pregnant mice.

Although the litter size and embryo number on D18 were unchanged in GDM mice (STZ) (***SI Appendix*, Fig. S1 *D* and *E***), birth weight was significantly reduced relative to non-GDM controls (Veh) (**Fig. 1*A***), indicating that this model recapitulates the SGA phenotype observed in human GDM. Further analysis revealed that both placentas and fetuses on D18 were significantly smaller in GDM pregnancies (**Fig. 1*B* and *C* and *SI Appendix*, Fig. S1*F***), suggesting that maternal hyperglycemia impairs placental growth and contributes to FGR. Notably, the fetal-to-placental weight ratio was significantly increased in GDM mice (**Fig. 1*D***), reflecting an elevated functional burden per unit of placental tissue. Hematoxylin and eosin (HE) staining demonstrated disrupted zonal differentiation in GDM placentas, including poorly defined boundaries between the junctional zone (JZ) and labyrinth zone (LZ) and aberrant extension of JZ cells into LZ regions (**Fig. 1*E***). Periodic acid Schiff (PAS) staining further showed abnormal accumulation of glycogen trophoblast (GlyT) islands within the LZ (**Fig. 1*F***), likely resulting from enhanced glycogen synthesis driven by excess glucose under hyperglycemia. Together, HE and PAS analyses indicate defective LZ development. In line with this, CD31 immunostaining demonstrated a significant reduction in capillary density within the LZ of GDM placentas (**Fig. 1 *G* and *H***), indicative of impaired angiogenesis. Since angiogenesis is closely coordinated with trophoblast syncytialization (26, 27), we next examined whether syncytial differentiation is altered by gestational hyperglycemia. In mice, MCT1 (*Slc16a1*) marks the maternal-facing syncytiotrophoblast layer I (SynT-I), whereas MCT4 (*Slc16a3*) marks the fetal-facing syncytiotrophoblast layer II (SynT-II)(28). Immunofluorescence staining revealed significantly increased expression of both MCT1 and MCT4 in GDM placentas (**Fig. 1*I***), indicating abnormal syncytialization. Given that MCT1/4 transport lactate, pyruvate, and ketone bodies(29), their elevated expression likely reflects a metabolic adaptation to maternal hyperglycemia.

**Fig. 1.**
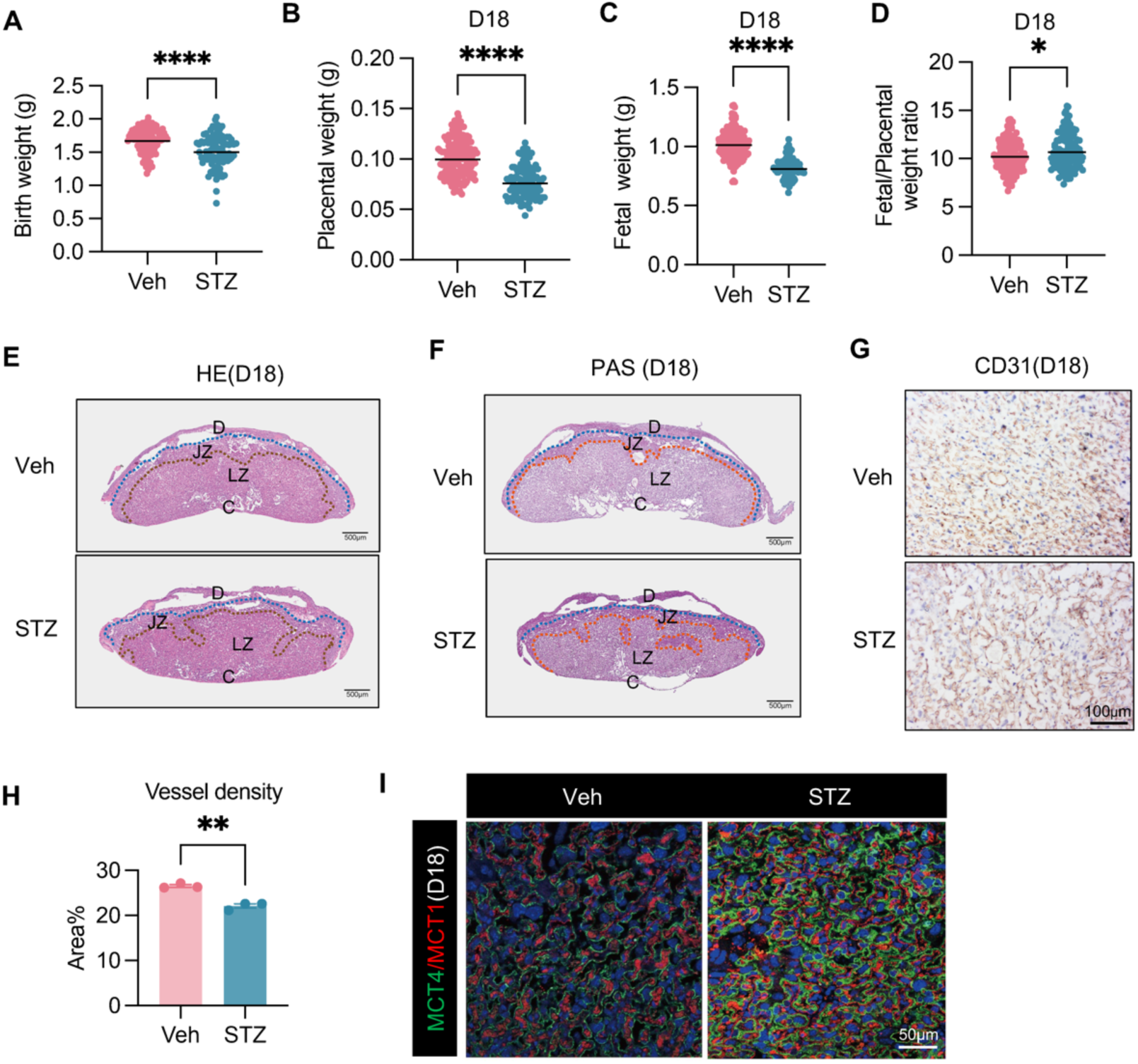
The gestational hyperglycemia induced by STZ in pregnant mice lead to abnormal placental structure and FGR. (*A*) Birth weight of the offspring in Veh (n=8) and STZ (n=9) pregnancies. Data represent the mean. **(*B-D*)** Placental weight (*B*), fetal weight (*C*), and fetal/ placental weight ratio (*D*) in Veh (n=11) or STZ (n=9) groups on D18. Data represent the mean. **(*E*)** Representative HE staining images of placental tissue sections in Veh and STZ placentas on D18 with dashed lines approximating the boundary between the decidual (D), junctional zone (JZ) and labyrinth zone (LZ). C represent Chorion. Scale bar, 500μm. **(*F*)** PAS staining for glycogen trophoblast. Scale bar, 500μm. **(*G*)** Representative CD31 immunostaining of the endothelium in Veh and STZ placental labyrinth zone on D18. Scale bar, 100μm. **(*H*)** Quantification of labyrinth endothelium density based on CD31 staining. Data represent the mean ± SEM. **(*I*)** Representative immunofluorescence staining of MCT1 and MCT4 in Veh and STZ placental labyrinth on D18. Scale bar, 50μm. In *A*, *B*, *C*, *D* and *H*, the analysis was carried out by two-tailed unpaired student’s t test. \**P* < 0.05, \*\**P* < 0.01, \*\*\*\**P* <0.0001.

Collectively, these findings demonstrate that maternal hyperglycemia disrupts multiple aspects of placental development, including growth, angiogenesis, syncytialization, and GlyT differentiation, thereby increasing the physiological burden on each unit of placental tissue. As a result, the compromised placenta fails to adequately support fetal growth, leading to FGR.

### Maternal hyperglycemia reprograms placental metabolic homeostasis in GDM by driving anabolic pathways

Given that maternal hyperglycemia induces aberrant expression of the syncytial/metabolic markers MCT1 and MCT4, along with abnormal accumulation of GlyT cells, we hypothesized that maternal hyperglycemia broadly disrupts metabolic homeostasis in the GDM placenta. To test this hypothesis, we performed untargeted metabolomic profiling of placentas from non-GDM control and GDM mice. In total, 1,039 metabolites were detected, of which 100 were significantly upregulated and 45 were downregulated in GDM placentas (***SI Appendix*, Fig. S2*A***). Metabolic pathway analysis revealed widespread metabolic perturbations (**Fig. 2*A***). Notably, the hexosamine biosynthetic pathway (HBP) was markedly elevated in GDM placentas, as indicated by increased levels of N-acetyl-D-glucosamine 6-phosphate (GlcNAc-6P) and uridine diphosphate N-acetylglucosamine (UDP-GlcNAc) (**Fig. 2*A* and *SI Appendix*, Fig. S2 *B* and *C***). Consistent with UDP-GlcNAc serving as a key substrate for protein O-glycosylation(30), GDM placentas exhibited increased levels of glycosylated proteins (***SI Appendix*, Fig. S2*D***). The accumulation of hexosamine intermediates and enhanced protein glycosylation indicate sustained exposure to a hyperglycemic environment, further validating the successful establishment of our GDM mouse model.

**Fig. 2.**
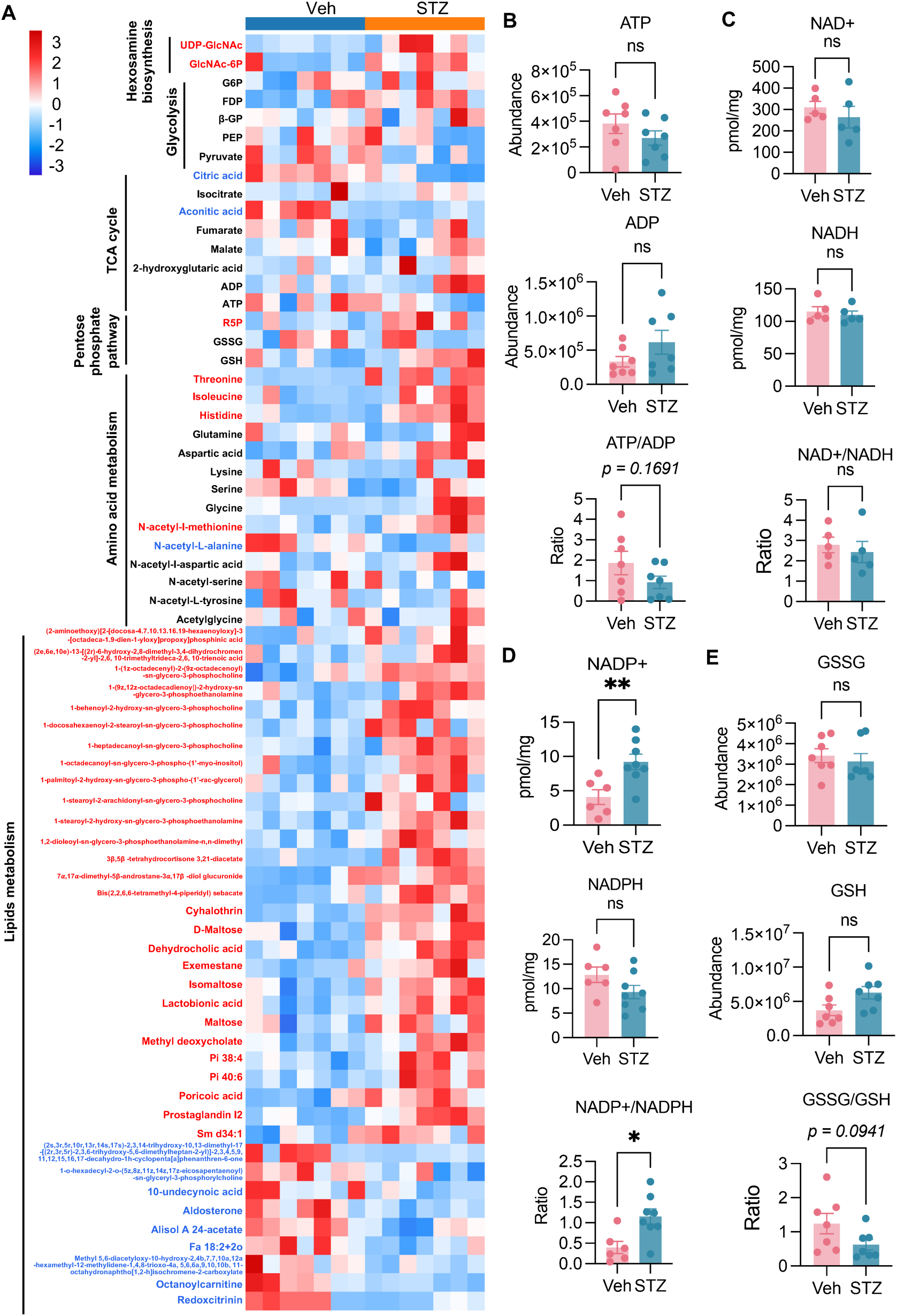
Placental tissues of GDM pregnant mice showed an increasing trend of anabolic metabolism. (*A*) Unsupervised heatmap of metabolomics analysis data of placental tissue on D18 from Veh (n=7) and STZ (n=7) groups detected by LC-MS. Rows are grouped by pathway (glycolysis, pentose phosphate pathway, TCA cycle, amino-acid metabolism, hexosamine biosynthesis, and lipid metabolism). **(*B*)** Relative abundance of ATP and ADP detected by LC-MS, and the ratio of ATP/ADP in Veh and STZ placentas on D18. **(*C*)** Quantification of NAD⁺, NADH and the NAD⁺/NADH ratio in Veh and STZ placentas on D18. (***D***) Quantification of NADP⁺, NADPH and the NADP⁺/NADPH ratio in Veh and STZ placentas on D18. **(*E*)** Relative abundance of oxidized glutathione (GSSG), reduced glutathione (GSH) detected by LC-MS, and the ratio of GSSG/GSH in Veh and STZ placentas on D18. In *C* and *E*, the analysis was carried out by two-tailed unpaired student’s t test. Data represent the mean ± SEM. ns, not significant. \**P* < 0.05, \*\**P* < 0.01.

Despite excessive glucose availability in GDM placentas, glycolytic intermediates were not significantly altered compared with non-GDM controls (**Fig. 2*A***). Most tricarboxylic acid (TCA) cycle metabolites also remained unchanged, with the exception of citrate and aconitic acid, which were significantly reduced in GDM placentas (**Fig. 2*A***), suggesting a potential suppression of TCA cycle activity. Absolute ATP levels tended to decrease in GDM placentas, whereas ADP levels were inversely increased (**Fig. 2 *A* and *B***). Although these opposing trends would be expected to lower the ATP/ADP ratio, the change did not reach statistical significance due to large variation among samples (**Fig. 2 *A* and *B***). Nevertheless, the downward trend suggests a compromised oxidative phosphorylation (OXPHOS). In contrast, NAD⁺ and NADH levels, as well as the NAD⁺/NADH ratio, were unchanged (**Fig. 2*C***), suggesting the engagement of compensatory mechanisms to preserve the redox balance. Interestingly, GDM placentas exhibited enhanced pentose phosphate pathway (PPP) activity, as evidenced by elevated ribose-5-phosphate (R5P) levels (**Fig. 2*A* and *SI Appendix*, Fig. S2*E***). Although the PPP is a major source of NADPH(31), we observed increased NADP⁺ levels and an elevated NADP⁺/NADPH ratio, accompanied by a decreased GSSG/GSH ratio (**Fig. 2 *D* and *E***), indicating heightened demand for reducing equivalents. Consistent with this, lipid biosynthesis was markedly enhanced, reflected by a substantial increase in lipid metabolites in GDM placentas (**Fig. 2A**). As NADPH is an essential reducing equivalent for anabolic reactions(32), its increased consumption by lipid synthesis likely contributes to the elevated NADP⁺ levels and NADP⁺/NADPH ratio, which may in turn further stimulate PPP flux. Among lipid-derived mediators, prostacyclin (PGI₂), a key placental lipid hormone, was significantly increased in GDM placentas (**Fig. 2*A* and *SI Appendix*, Fig. S2*F***). Because PGI₂ is predominantly produced by syncytiotrophoblasts during placenta development(33–35), its elevation may reflect dysregulated placental syncytialization, consistent with increased expression of syncytial markers, including MCT1 and MCT4 (**Fig. 1*I***). We also observed elevated γ-aminobutyric acid (GABA) levels in GDM placentas (***SI Appendix*, Fig. S2*G***). Given that glutamine levels were unchanged (***SI Appendix*, Fig. S2*H***), the increased GABA is likely derived from glucose-driven metabolic flux through α-ketoglutarate and glutamate. As both PGI₂ and GABA have been implicated in promoting vascularization(27, 36, 37), their upregulation may represent compensatory responses to impaired angiogenesis in GDM placentas. In addition, multiple amino acids were elevated in GDM placentas (**Fig. 2*A***), potentially reflecting enhanced glucose-driven amino acid biosynthesis. Disaccharides, including maltose and lactose, as well as the oligosaccharide lacto-N-tetraose (LNT), were also significantly increased (***SI Appendix*, Fig. S2 *I-K***), consistent with previous reports that maternal hyperglycemia promotes lactogenesis(38, 39).

Collectively, these findings demonstrate that gestational hyperglycemia profoundly reprograms placental metabolism by activating multiple anabolic pathways—including the HBP, the PPP, lipid synthesis, amino acid production, and lactogenesis—while concurrently suppressing the TCA cycle and OXPHOS. This coordinated metabolic rewiring underscores the substantial anabolic burden and bioenergetic stress imposed on the GDM placenta.

### Hyperglycemia-driven metabolic reprogramming remodels the epigenetic landscape via acetyl-CoA-mediated histone hyperacetylation in GDM placentas

Our untargeted metabolomic analysis revealed that maternal hyperglycemia comprehensively reprograms placental metabolism in GDM mice (**Fig. 2**). This shift is characterized by the promotion of the HBP and lipid biosynthesis, alongside the concomitant suppression of the TCA cycle and OXPHOS. Because both HBP and lipid biosynthesis require acetyl-CoA as a critical metabolic intermediate(40, 41), we hypothesized that glucose-derived citrate is redirected from the mitochondria into the cytosol to fuel acetyl-CoA production for anabolism (**Fig. 3*A***). In support of this hypothesis, we observed an inverse relationship between citrate and acetyl-CoA levels in GDM placentas compared to non-GDM controls: citrate levels were markedly decreased, while acetyl-CoA levels were significantly elevated (**Fig. 2*A* and 3 *B* and *C***). This reciprocal pattern suggests an accelerated conversion of citrate into acetyl-CoA within GDM placental tissues. This enhanced acetyl-CoA pool likely drives the observed increase in lipid biosynthesis, a finding supported by both our metabolomic data and intensified lipid droplet staining (**Fig. 2*A* and 3*D* and *SI Appendix*, Fig. S3*A***). Beyond its role as a substrate for anabolism, acetyl-CoA is a well-established regulator of histone acetylation, a central mechanism of epigenetic control(42, 43). We reasoned that the diversion of carbon flux toward acetyl-CoA might also alter histone acetylation dynamics. Such metabolic-epigenetic coupling could underlie the extensive epigenetic reprogramming that influences placental development in GDM. To test this, we assessed global histone acetylation and found that total H3 acetylation, H3K27ac, H3K9ac, H3K14ac, and H3K18ac were significantly elevated in GDM placentas (**Fig. 3*E*, *SI Appendix*, Fig. S3*B***). Notably, H3K27ac showed the most pronounced change, increasing approximately 3-fold in GDM placentas compared to controls (**Fig. 3*E*, *SI Appendix*, Fig. S3*B***). Immunofluorescence staining further confirmed increased nuclear H3K27ac levels within the labyrinth trophoblast cells of D18 GDM placentas (**Fig. 3*F***).

**Fig. 3.**
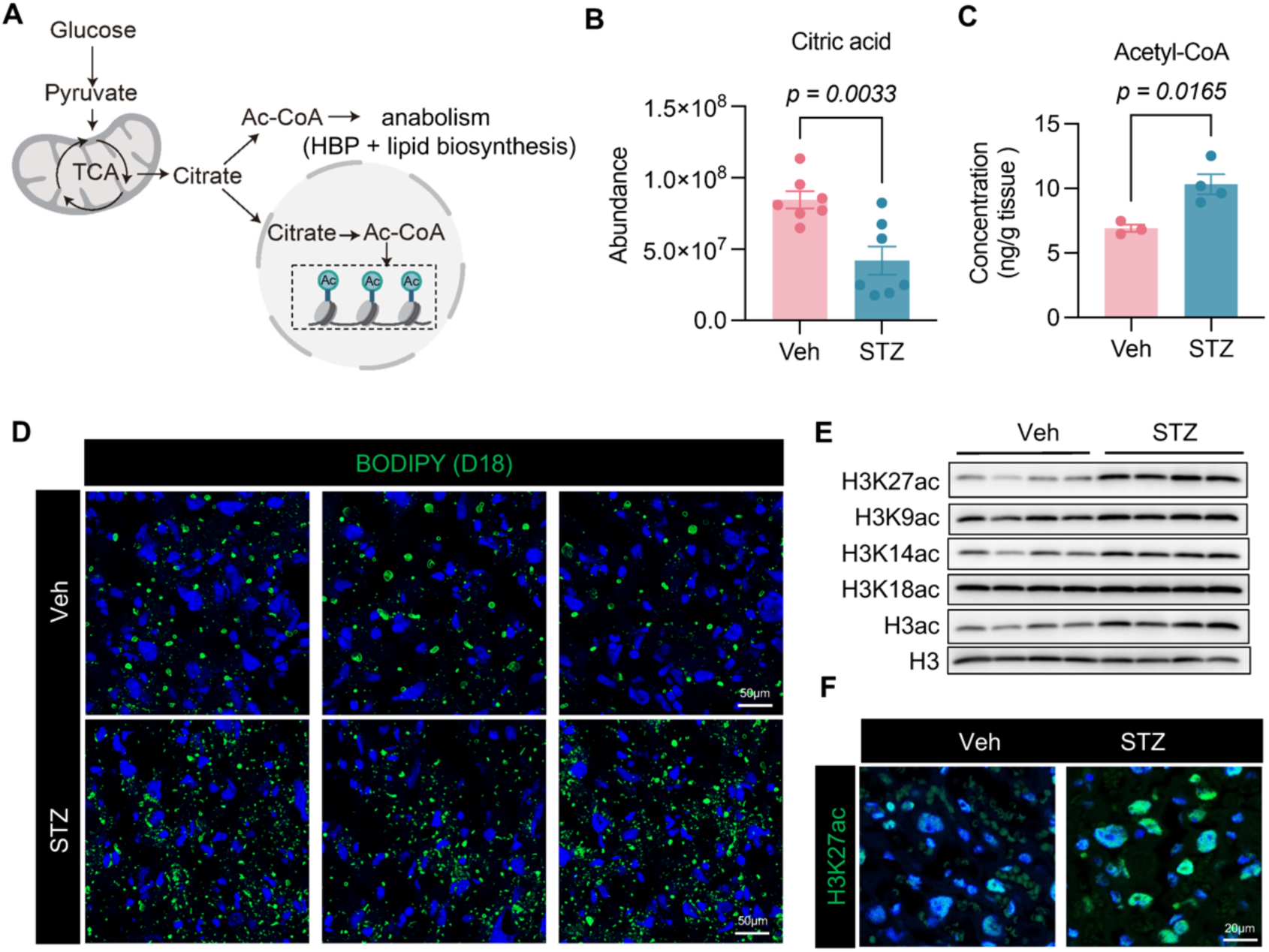
The placental tissues of GDM mice showed an increase in lipid droplet accumulation and histone acetylation. (***A***) Schematic illustrating glucose-derived citrate export and ACLY-dependent acetyl-CoA production linking metabolism to histone acetylation. (***B***) Relative abundance of citric acid in Veh and STZ placentas on D18 detected by LC-MS. (***C***) Quantification of Acetyl-CoA concentration in Veh and STZ placentas on D18. Data represent the mean ± SEM. Two-tailed unpaired Student’s t-test. *P*=0.0165. (***D***) Representative BODIPY staining in Veh and STZ placentas on D18. Scale bar, 50μm. (***E***) Immunoblotting analysis of histone acetylation marks (H3K27ac, H3K9ac, H3K14ac, H3K18ac, and total H3ac) in Veh and STZ placentas on D18 with total H3 as a loading control. (***F***) Representative immunofluorescence staining of H3K27ac in Veh and STZ placentas on D18. Scale bar, 20μm.

Taken together, these findings demonstrate that hyperglycemia-driven metabolic reprogramming redirects glucose-derived citrate toward acetyl-CoA production. This metabolic shift promotes histone hyperacetylation, thereby remodeling the placental epigenetic state.

### Histone hyperacetylation-associated epigenetic remodeling alters the placental transcriptomic landscape in GDM pregnancies

Since epigenetic remodeling is directly linked to the regulation of gene expression(44, 45), we next performed RNA sequencing (RNA-seq) to determine whether gestational hyperglycemia disrupts the transcriptional profile in the placentas of GDM mice compared to non-GDM controls. Principal component analysis (PCA) revealed a clear separation between GDM and non-GDM placentas (***SI Appendix*, Fig. S4*A***), indicating a profoundly altered transcriptional landscape in response to maternal hyperglycemia. Differential expression analysis identified 1,974 differentially expressed genes (DEGs), including 1,401 upregulated and 573 downregulated genes (**Fig. 4*A***). Notably, the number of upregulated genes was approximately 2.44-fold higher than that of downregulated genes (**Fig. 4*B***). This global bias toward transcriptional activation is consistent with the well-established concept that histone hyperacetylation—particularly increased H3K27ac (**Fig. 3 *E* and 3*F***)—generally promotes gene transcription(46, 47). Interestingly, genes encoding histone acetyltransferases (HATs) were consistently upregulated, whereas genes encoding histone deacetylases (HDACs) were downregulated (**Fig. 4*C***), suggesting that increased histone acetylation in the GDM placenta is driven by the synergistic effect of enhanced acetyl group addition and reduced removal. This coordinated expression pattern of histone acetyltransferases and deacetylases likely reflects an adaptive response to elevated acetyl-CoA levels in the GDM placenta.

**Fig. 4.**
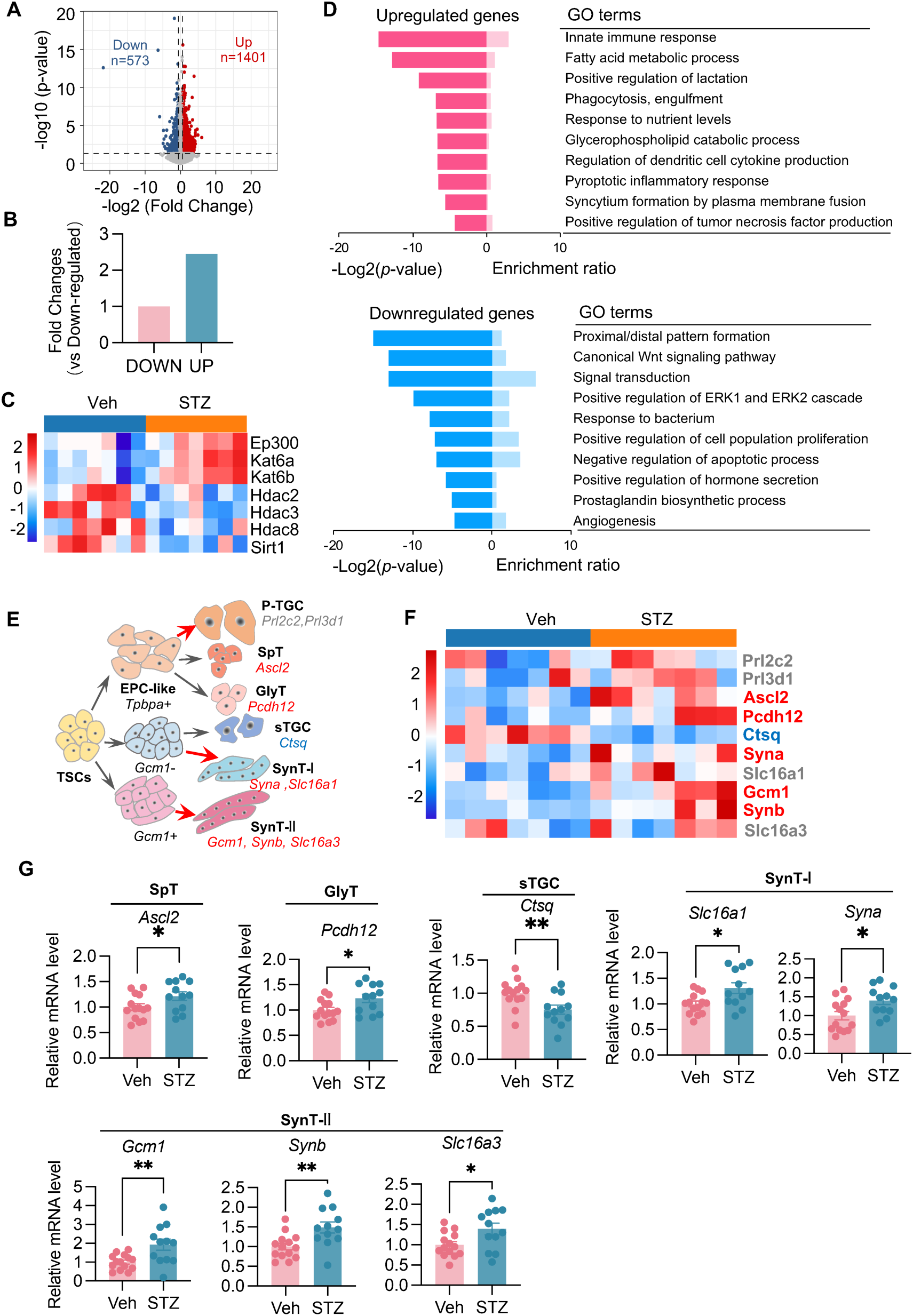
RNA-seq showed global transcriptional activation and trophoblast lineage imbalance in GDM placenta. (***A***) Volcano plot of different expressed genes (DEGs) between Veh and STZ placentas on D18. Dashed vertical lines mark the fold-change threshold and the horizontal line marks the significance cutoff. (***B***) The relative differences in the number of DEGs between Veh and STZ placentas on D18. (***C***) Heatmap showing expression changes of genes encoding histone acetyltransferases (HATs) and histone deacetylases (HDACs) in Veh and STZ placentas on D18. **(*D*)** GO analysis of upregulated and downregulated DEGs. (***E***) Schematic of trophoblast lineage specification and representative markers for SpT, GlyT, sTGC, SynT-I, and SynT-II. **(*F*)** Heatmap of representative trophoblast lineage marker expression in Veh and STZ placentas on D18. **(*G*)** RT-qPCR analysis of trophoblast lineage markers. The values are normalized to *Actb* and indicated as the mean ± SEM. Two-tailed unpaired Student’s t-test. \**P* <0.05, \*\**P* <0.01.

Gene Ontology (GO) analysis of the DEGs revealed that upregulated genes were significantly enriched in pathways related to innate immunity, inflammatory responses, metabolism, and syncytialization (**Fig. 4*D***). In contrast, downregulated genes were enriched in pathways associated with proximal/distal pattern formation, Wnt and ERK signaling, antibacterial defense, positive regulation of cell proliferation, negative regulation of apoptosis, and angiogenesis (**Fig. 4*D***). These transcriptional changes align with our observations of enhanced syncytialization, impaired angiogenesis, and altered metabolism in the GDM placenta (**Fig. 1 *G*–*I* and Fig. 2**). Furthermore, we also uncover potentially heightened inflammation and apoptosis, alongside compromised cell proliferation and antimicrobial defense. Collectively, these transcriptional alterations may contribute to the reduced placental size and impaired fetal growth observed in GDM mice (**Fig. 1 *A*–*D* and *SI Appendix*, Fig. S1*F***).

In addition to GO analysis, Gene Set Enrichment Analysis (GSEA) using curated datasets for trophoblast differentiation demonstrated a general trend toward increased expression of trophoblast differentiation genes (***SI Appendix*, Fig. S4*B***). To further characterize trophoblast lineage specification, we analyzed key trophoblast lineage markers and observed significantly increased expression of markers for spongiotrophoblasts (SpT), GlyT, SynT-I and SynT-II (**Fig. 4 *E* and *F* and *SI Appendix*, Fig. S4*C***). These findings are consistent with the abnormal JZ, LZ, SynT, and GlyT structures observed in GDM placentas (**Fig. 1**). Interestingly, while immunostaining revealed markedly increased protein levels of MCT1 (*Slc16a1*) and MCT4 (*Slc16a3*), only modest increases in their respective mRNA levels were detected (**Fig. 1*G* and Fig. 4*F* and *SI Appendix*, Fig. S4*C***), suggesting the involvement of post-transcriptional mechanisms that enhance protein stability. Notably, *Ctsq*, a marker of sinusoidal trophoblast giant cells (sTGCs), was significantly downregulated (**Fig. 4 *E* and *F* and *SI Appendix*, Fig. S4*C***), indicating impaired sTGC differentiation. Because trophoblast giant cells are reported to play critical roles in host defense against microbes, including bacteria (48), we speculate that the reduced expression of antibacterial defense genes observed in GDM placentas may be attributable, at least in part, to defective sTGC differentiation (**Fig. 4 *D* and 4*F***). These aberrant expression patterns of trophoblast lineage markers were further validated via qPCR (**Fig. 4*G***).

In summary, these data demonstrate that hyperglycemia-driven histone hyperacetylation remodels the epigenome during placental development, globally reshaping gene expression by preferentially promoting transcriptional activation. This widespread transcriptional imbalance disrupts placental development and function in GDM pregnancies, ultimately contributing to FGR and SGA outcomes.

### ACLY upregulation in the nucleus and cytosol represents a key placental metabolic adaptation to maternal hyperglycemia

GO analysis revealed that a subset of genes upregulated in GDM placentas was significantly enriched in multiple metabolic pathways, including lipid metabolic processes (**Fig. 4*D***), suggesting that GDM placentas adopt a metabolic adaptation-related gene expression program in response to maternal hyperglycemia. To further investigate this adaptation, we performed GSEA focusing on specific metabolic pathways. Although the enrichment did not reach statistical significance, we observed a trend toward increased expression of genes associated with the PPP and glycogen synthesis (***SI Appendix*, Fig. S5 *A* and *B***), consistent with our metabolomic analyses showing increased levels of PPP metabolites and elevated GlyT (**Figs. 1*F* and 2*A***). Notably, we also observed a trend toward increased expression of TCA cycle genes that did not reach statistical significance (***SI Appendix*, Fig. S5C**), whereas genes related to OXPHOS were significantly downregulated (**Fig. 5*A* and *SI Appendix*, Fig. S5*D***). Consistent with this, our untargeted metabolomic data showed reduced levels of TCA cycle metabolites and decreased ATP production (**Figs. 2 *A* and *B***), suggesting that suppression of both TCA cycle activity and OXPHOS is primarily driven by downregulation of OXPHOS genes. Finally, genes associated with lipid biosynthesis were significantly upregulated in GDM placentas compared with non-GDM controls (**Fig. 5*B* and *SI Appendix*, Fig. S5*E***), consistent with the marked increase in lipid metabolites and lipid droplet formation observed in GDM placentas (**Figs. 2*A* and 3*D***). Together, these metabolic gene expression patterns in GDM placentas are consistent with our untargeted metabolomic data and reflect metabolic adaptation to maternal hyperglycemia at the transcriptional level.

**Fig. 5.**
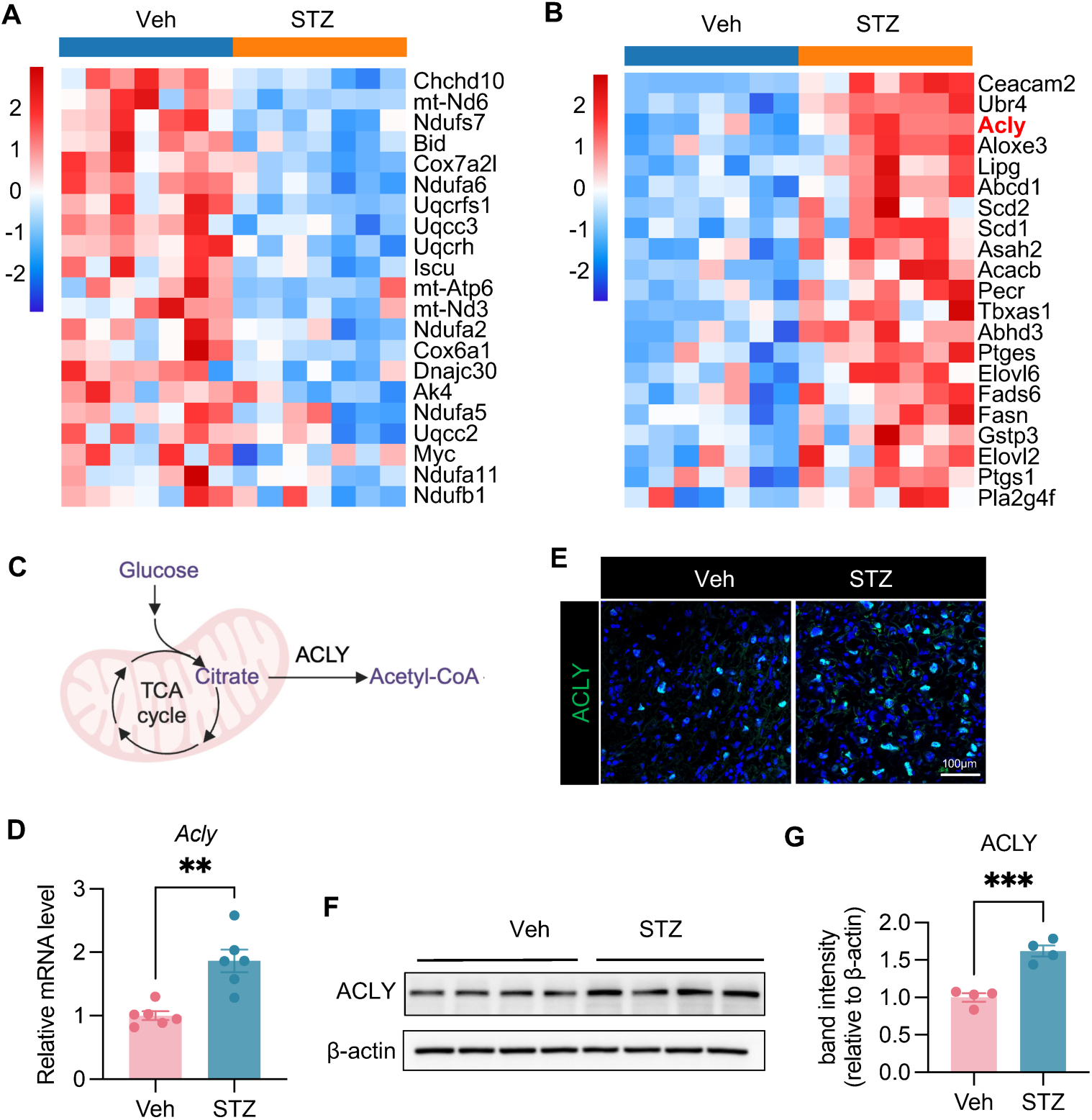
Transcriptome highlight suppressed OXPHOS and increased FA biosynthesis accompanied by ACLY induction in GDM placenta. (*A*) Heatmap of representative OXPHOS related genes in Veh and STZ placentas on D18. (***B***) Heatmap of representative FA biosynthesis related genes in Veh and STZ placentas on D18. (***C***) Schematic of citrate-to-acetyl-CoA conversion by ACLY. (***D***) RT-qPCR analysis of *Acly* in Veh and STZ placentas on D18. The values are normalized to *Actb* and indicated as the mean ± SEM. \*\**P* <0.01. **(*E*)** Representative immunofluorescence staining of ACLY in Veh and STZ placentas on D18. Scale bars, 100μm. (***F***) Immunoblotting analysis of ACLY in Veh and STZ placentas on D18 with β-actin as a loading control. **(*G*)** Densitometric quantification of ACLY normalized to β-actin in Veh and STZ placentas on D18. Data represent the mean ± SEM. Two-tailed unpaired Student’s t-test, \*\*\**P* < 0.001.

Our data support suppression of OXPHOS and activation of acetyl-CoA production to fuel histone acetylation and anabolic processes, including lipid biosynthesis and the HBP. We therefore proposed that glucose-derived carbons may be redirected toward acetyl-CoA production. To investigate this possibility, we analyzed our transcriptomic data in detail and found that ATP citrate lyase (ACLY), a rate-limiting enzyme that converts citrate into cytosolic and nuclear acetyl-CoA(49–52), was significantly upregulated (**Fig. 5 *B* and *C* and *SI Appendix*, Fig. S5*F***). qPCR further confirmed the transcriptomic results, showing that *Acly* mRNA levels were significantly elevated in GDM placentas compared with non-GDM controls (**Fig. 5*D***). Immunofluorescence staining revealed that ACLY is predominantly expressed in syncytiotrophoblasts (giant nuclei) in non-GDM placentas (**Fig. 5*E***). In contrast, both mononucleated (small nuclei) and multinucleated (giant nuclei) trophoblast cells in GDM placentas exhibited markedly elevated ACLY expression, likely reflecting adaptation to chronic hyperglycemic stress (**Fig. 5*E***). Notably, ACLY displayed strong nuclear localization in syncytiotrophoblasts (**Fig. 5*E***), consistent with the significantly increased H3K27ac levels observed in these cells in GDM placentas (**Fig. 3*F***). Given that nuclear ACLY has been reported to promote H3K27 acetylation and epigenetically regulate gene expression (52), we propose that enhanced nuclear ACLY in syncytiotrophoblasts of GDM placentas supplies nuclear acetyl-CoA to support epigenetic remodeling. Western blot analysis further confirmed a significant increase in ACLY protein abundance in GDM placentas (**Fig. 5 *F* and *G***).

Collectively, these data indicate that the placenta metabolically adapts to maternal hyperglycemia through reprogramming of metabolic gene expression. Among these changes, upregulation of ACLY in both the nucleus and cytosol appears to facilitate rapid redirection of glucose-derived carbons toward acetyl-CoA production in these compartments, thereby supporting both nuclear histone hyperacetylation and cytosolic acetyl-CoA–dependent anabolic processes.

### ACLY upregulation and nuclear localization, increased acetyl-CoA, and enhanced histone acetylation are conserved features of human GDM placentas

To further investigate how gestational hyperglycemia affects placental and fetal development in patients with GDM, we collected clinical data from 146 volunteer participants at Hangzhou Women’s Hospital. Among these participants, 89 were diagnosed with GDM, and the remaining 57 served as non-GDM controls. To minimize potential confounding effects from comorbidities, advanced maternal age, and pre-pregnancy obesity, we excluded participants with additional pregnancy complications, those aged ≥35 years, and those with a pre-pregnancy body mass index (BMI) ≥28 kg/m². Because our primary objective was to examine the effects of maternal hyperglycemia on placental development and fetal health, we further excluded GDM participants with HbA1c <5.5% after dietary and exercise management (53). Based on these criteria, we ultimately included 52 non-GDM controls and 32 GDM cases that were comparable in overall health status, maternal age, and pre-pregnancy body mass index (BMI) (**Fig. 6*A***). The GDM diagnosis was further confirmed by significantly elevated HbA1c levels and increased oral glucose tolerance test (OGTT) glucose concentrations at 0, 1, and 2 hours after glucose load in the GDM group compared with controls (**Fig. 6*A***). Although overall neonatal birth weight was comparable between the two groups, 4 of 32 (12.5%) pregnancies in the GDM group resulted in SGA neonates (**Fig. 6 *A*–*C***), which is comparable with previous reports in Asian populations showing SGA incidences in GDM pregnancies of approximately 13.26%(15). In contrast to the well-established association between GDM and LGA infants(4), we observed only a modest, non-significant increase in the incidence of LGA neonates in the GDM group compared with controls (**Fig. 6 *A* and *D***). This may be because the GDM cases included in our study excluded the confounding effect of pre-pregnancy obesity. In addition, neonatal intensive care unit (NICU) admission rates were approximately 4-fold higher among neonates born to GDM mothers than among non-GDM control neonates (**Fig. 6 *A* and *E***), consistent with prior studies(54). Collectively, these findings indicate that our patient cohort is representative and comparable to previously reported GDM populations. However, we were unable to evaluate the relationship between placental growth and fetal growth because accurate placental weight data were unavailable due to incomplete clinical records.

**Fig. 6.**
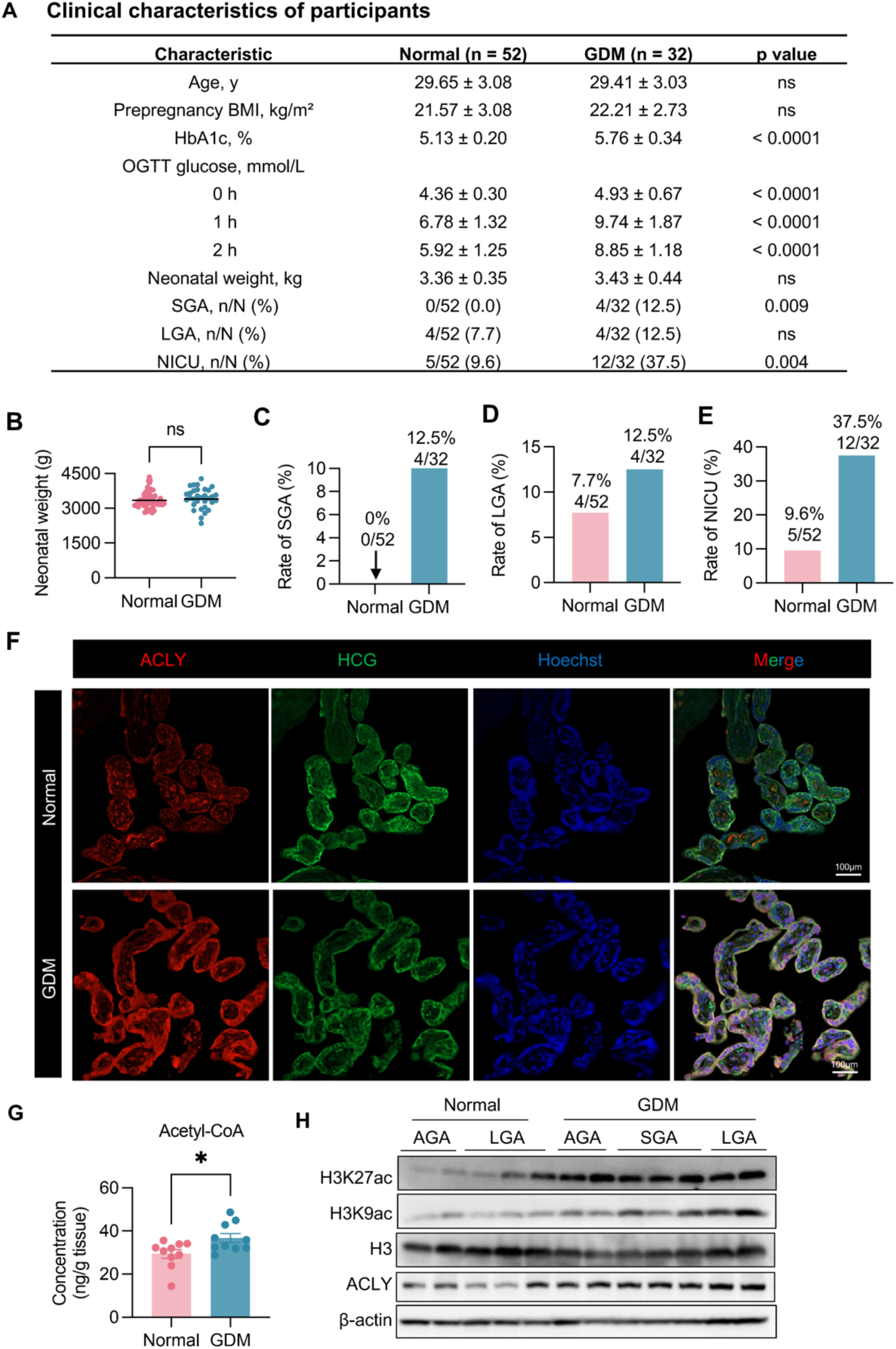
Clinical cohort characterization and elevated placental ACLY and H3K27ac/ H3K9ac in GDM. (*A*) Clinical characteristics of participants including age, pre-pregnancy BMI, HbA1c, OGTT (0 h, 1 h, 2 h), neonatal birth weight, SGA, LGA, and NICU in normal and GDM groups. Values are mean ± SD or n/N (%). Continuous variables were compared using unpaired two-tailed t tests. Categorical variables were compared using Fisher’s exact tests. ns, not significant. **(*B*)** Neonatal birth weight in normal and GDM groups. Data represent the mean. Two-tailed unpaired Student’s t-test. ns, not significant. **(*C–E*)** Rates of SGA (*C*), LGA (*D*), and NICU admission (*E*) in normal and GDM groups, shown as n/N (%). **(*F*)** Representative immunofluorescence images of ACLY and hCG with nuclei counterstained by Hoechst in normal and GDM term placental villis. Scale bar, 100μm. **(*G*)** Quantification of acetyl-CoA concentration in term placental tissues from normal and GDM. Data represent the mean ± SEM. Two-tailed unpaired Student’s t-test. \**P* < 0.05. **(*H*)** Representative immunoblots of H3K27ac/ H3K9ac and ACLY in term placental tissues from normal and GDM. Total H3 and β-actin serve as loading controls.

Based on findings from our GDM mouse model demonstrating that maternal hyperglycemia disrupts acetyl-CoA homeostasis through upregulation of ACLY, which further induces histone hyperacetylation-related epigenetic remodeling during placental development, we next examined ACLY expression and subcellular localization in human placentas. Term placental villous tissues were collected, and human chorionic gonadotropin (hCG), a canonical marker of syncytiotrophoblasts and a major trophoblast-derived hormone, was used to delineate the syncytiotrophoblast layer covering the villi(55). Immunofluorescence staining revealed that term human placental villi were predominantly composed of syncytiotrophoblasts (**Fig. 6*F***). In non-GDM control placentas, ACLY was primarily localized to the cytosol of syncytiotrophoblasts, whereas GDM placentas exhibited markedly elevated ACLY levels in both the cytosol and nucleus (**Fig. 6*F***), consistent with our observations in the GDM mouse model. Consistently, acetyl-CoA levels in term placental tissues from GDM patients were significantly higher than those in non-GDM controls (**Fig. 6*G***). We next examined whether classic histone acetylation, such as H3K27ac and H3K9ac, were similarly altered in placentas from human GDM patients. Term placental tissues were collected from two non-GDM controls with AGA neonates, three non-GDM controls with LGA neonates, two GDM patients with AGA neonates, three GDM patients with SGA neonates, and two GDM patients with LGA neonates. Western blot analysis showed that placentas from GDM patients, regardless of birth outcomes, consistently exhibited higher levels of H3K27ac and H3K9ac. In contrast, placentas from non-GDM controls with different birth outcomes showed some variability but overall substantially lower levels of histone acetylation (**Fig. 6*H* and *SI Appendix*, Fig. S6 *A* and *B***). Interestingly, ACLY protein levels in these placentas closely paralleled the histone acetylation status (**Fig. 6*H* and *SI Appendix*, Fig. S6*C***), suggesting that ACLY-dependent acetyl-CoA production is associated with histone hyperacetylation in human GDM placentas.

These data indicate that maternal hyperglycemia in GDM pregnancies is associated with an increased risk of SGA outcomes and NICU admission in this Chinese cohort. Notably, all GDM placentas exhibited conserved upregulation and nuclear localization of ACLY, increased acetyl-CoA levels, and enhanced histone acetylation compared with non-GDM placentas, consistent with observations from our GDM mouse model. Collectively, we propose that aberrant activation of the ACLY-acetyl-CoA-histone hyperacetylation axis represents a conserved placental response to maternal hyperglycemia and may contribute to placental dysfunction and heterogeneous birth outcomes in GDM pregnancies.

## Discussion

GDM is characterized by maternal hyperglycemia and is therefore considered a major metabolic complication of pregnancy (1, 2). Exposure to maternal hyperglycemia poses substantial challenges to fetal development (3–5). One direct consequence of GDM is an increased incidence of abnormal birth weight compared with non-GDM pregnancies, encompassing both LGA and SGA infants (3–5). Previous study has reported an average LGA incidence of 16.3% in GDM pregnancies (range, 3.5%–37.7%), corresponding to a 1.66-fold increase relative to non-GDM pregnancies(54). Consistent with these reports, our clinical data demonstrate a 1.62-fold increased risk of LGA, with an incidence of 12.5% in GDM pregnancies compared with 7.7% in non-GDM pregnancies. In contrast, SGA outcomes have generally been reported to be less frequent in GDM pregnancies, with an average incidence of approximately 7% (56). However, the incidence of GDM-associated SGA appears to be highly dependent on race and ethnicity. For example, Asian populations exhibit a higher incidence approximately at 13.26%(15). In our cohort from a Chinese population, we observed a similar SGA incidence of 12.25% in GDM pregnancies, whereas no SGA cases were detected in non-GDM pregnancies. These findings suggest that maternal hyperglycemia may act as a population-specific risk factor for SGA in GDM pregnancies. The association between GDM and LGA is relatively intuitive, as excessive maternal glucose availability promotes fetal overgrowth. Although placentas from GDM pregnancies often exhibit structural abnormalities and functional insufficiency, GDM-associated LGA pregnancies typically display a reduced fetus-to-placenta weight ratio, indicating that placental overgrowth exceeds fetal overgrowth (7). This observation suggests that fetal overgrowth is supported by placental enlargement and enhanced transplacental nutrient transfer, including glucose, lactogens, free fatty acids, and amino acids. In contrast, SGA is typically associated with FGR, making it paradoxical that excessive nutrient availability in GDM pregnancies could result in impaired fetal growth. To investigate this paradox, we established a GDM mouse model that recapitulates the SGA phenotype observed in human GDM pregnancies. In this model, both placental and fetal weights were significantly reduced in GDM mice compared with non-GDM controls, accompanied by an increased fetus-to-placenta weight ratio. This pattern is opposite to that observed in human GDM-associated LGA cases, in which placental overgrowth predominates. Despite evident structural abnormalities and impaired vacuolation in SGA-associated GDM placentas in mice, the elevated fetus-to-placenta weight ratio indicates that increased nutrient availability per unit placental mass may partially compensate for placental insufficiency, enabling relatively greater fetal growth relative to placental size. Together, these findings suggest that GDM-associated SGA primarily arises from placental underdevelopment–driven FGR rather than insufficient nutrient supply.

As a metabolic disorder, GDM-associated hyperglycemia produces heterogeneous fetal outcomes, ranging from overgrowth (LGA) to growth restriction (SGA), likely reflecting differences in placental metabolic adaptation and maternal–fetal nutrient partitioning. To explore the metabolic basis of these divergent outcomes, we performed comprehensive untargeted metabolomic analyses of placentas from GDM and non-GDM pregnant mice. We observed significantly elevated anabolic metabolic activity in SGA-associated mouse GDM placentas compared with non-GDM controls, including increased flux through the HBP, the PPP, lipid and fatty acid synthesis, lactogenesis, and amino acid biosynthesis. This enhanced anabolic activity is consistent with previous studies in human GDM placental samples and in vitro models (19, 20, 57), suggesting a conserved metabolic adaptation to chronic glucose overload in placentas under hyperglycemic conditions in both mouse models and human GDM pregnancies. Notably, our study is the first to reveal that the TCA cycle and OXPHOS are suppressed in the GDM mouse placenta despite excessive glucose availability, indicating preferential diversion of glucose toward anabolic biosynthesis rather than mitochondrial energy production. Collectively, these data suggest that enhanced anabolism and sufficient nutrient availability are common features of GDM placentas and therefore are unlikely to account for the heterogeneous birth outcomes.

A large body of literature has established that metabolic reprogramming can profoundly reshape the epigenetic landscape through mechanisms ranging from supplying metabolic intermediates for epigenetic modifications to regulating the activity of epigenetic enzymes(58–61). Indeed, several studies have identified epigenetic alterations in GDM placentas compared with non-GDM placentas, including changes in histone acetylation, histone methylation, and DNA methylation(22, 23, 62). Our analyses revealed elevated levels of acetyl-CoA, a key metabolic intermediate that serves as a substrate for both anabolic metabolism and histone acetylation (40–43), as a prominent feature of the placental response to maternal hyperglycemia. Proper acetyl-CoA metabolism is essential for maintaining histone acetylation during syncytialization in human placental cells(17, 18). However, the impact of elevated acetyl-CoA production under hyperglycemic conditions on histone acetylation and placental development remains poorly understood. Here, we demonstrate that elevated acetyl-CoA levels in both human and mouse GDM placentas are associated with histone hyperacetylation, including increased H3K27ac and H3K9ac. Interestingly, a previous study reported reduced H3K9ac levels in human GDM placentas(23), whereas we observed the opposite trend in both human and mouse GDM placentas. Whether this discrepancy reflects population-specific differences remains unclear. Nevertheless, it is widely accepted that GDM placentas exhibit enhanced acetyl-CoA–driven anabolic metabolism, including lipid biosynthesis and activation of the HBP(19, 20, 57), supporting increased acetyl-CoA availability in GDM placentas. Elevated acetyl-CoA levels have been shown to promote histone hyperacetylation in both pathological and physiological contexts, including cancer and postimplantation embryonic development (43, 63). Therefore, the mechanism underlying reduced H3K9ac in certain GDM populations remains difficult to reconcile with the high acetyl-CoA availability observed under maternal hyperglycemia.

Consistent with previous studies showing that histone hyperacetylation generally promotes gene transcription (46, 47), we observed 2.44-fold more upregulated genes than downregulated genes in GDM mouse placentas. Aberrant upregulation of genes associated with innate immunity and inflammatory responses suggests an elevated inflammatory state in GDM placentas. Because placental inflammation is widely considered an important contributor to FGR and SGA(16), histone hyperacetylation–driven activation of inflammatory gene expression may represent a potential mechanism underlying GDM-associated FGR and SGA. In addition, the upregulation of syncytialization-related genes and the downregulation of genes involved in angiogenesis, anti-apoptotic pathways, and cellular proliferation indicate transcriptional dysregulation contributing to placental underdevelopment in GDM. Together, these findings suggest that histone hyperacetylation–driven transcriptional alterations may represent a key mechanism contributing to placental dysfunction and impaired placental development in GDM, ultimately leading to FGR and SGA. Interestingly, our data further indicate that metabolic genes are broadly reprogrammed in GDM placentas, with increased expression of lipid synthesis genes and decreased expression of OXPHOS-related genes. The expression patterns of these metabolic genes closely mirror the metabolic features observed in our metabolomic analyses, suggesting coordinated metabolic adaptation during placental development. Among these metabolic genes, ACLY—a rate-limiting enzyme that converts glucose-derived citrate to acetyl-CoA (49–52)—is upregulated in both the cytosol and nucleus of syncytiotrophoblasts in GDM placentas. Notably, the upregulation and nuclear localization of ACLY, enhanced acetyl-CoA production, increased lipid biosynthesis, and histone hyperacetylation are conserved between mouse and human GDM placentas. These findings suggest the existence of a conserved ACLY–acetyl-CoA axis that links hyperglycemia to anabolic metabolism and histone hyperacetylation–mediated epigenetic regulation.

Given that excessive nutrient availability through anabolic metabolism is a common feature of GDM placentas regardless of fetal growth outcome, we propose that heterogeneous fetal outcomes in GDM arise not from differences in global nutrient availability or global histone acetylation levels, but from histone hyperacetylation–driven transcriptional alterations that affect placental development.

In summary, our analyses of both mouse models and human GDM placentas reveal that the ACLY–acetyl-CoA metabolic pathway plays a central role in placental adaptation to maternal hyperglycemia, coordinating anabolic metabolism and epigenetic remodeling to influence placental development and fetal growth outcomes. This study provides new mechanistic insight into how GDM-associated hyperglycemia impacts placental function and fetal development. Future studies will investigate whether targeting ACLY may represent a potential therapeutic strategy to prevent abnormal fetal growth in GDM pregnancies.

## Data availability

The sequencing data generated in this study have been deposited in the Gene Expression Omnibus database under accession code GSE316949. Source data are provided with this paper.

## Acknowledgements

This work was supported by the National Natural Science Foundation of China (82271652 to XN.L., 82502018 to MY.L.), the Natural Science Foundation Key Program of Zhejiang Province (LZ26H040002 to XN.L.), the Construction Fund of Key Medical Disciplines of Hangzhou (2025HZPY08), the Medicine and Health Science and Technology Project of Zhejiang Province (2024KY1116 to MY.L.).

## Author contributions

The project was initiated and supervised by MY.L. and XN.L. The experiments were designed by MY.L. and B.H., with MY.L., KE.J., SH.Q. performing the experiments. DD.C., WN.X. and YC.H. helped to perform part of the experiments. SH.Q., CH.W., HY.W. and YW.L. contributed to the collection of clinical samples. MY.L., B.H. providing the bioinformatic analyses. MY.L., B.H. and XN.L. participated in the quality control of the study. The manuscript was written by MY.L., B.H. and XN.L. All authors participated in the discussion, revision, and approval of the manuscript.

## Materials and Methods

### Clinical data and human placental sample collection

This observational study was conducted in Hangzhou, Zhejiang, China. A total of 89 pregnant women diagnosed with GDM and 57 pregnant women without GDM were initially recruited in 2025. The study protocol was approved by the Ethics Committee of Hangzhou Women’s Hospital (Ethic No. 2025-69) and was performed in accordance with the Declaration of Helsinki. Written informed consent was obtained from all participants prior to enrollment. GDM was diagnosed at 24–28 weeks of gestation using a 75g OGTT according to the International Association of the Diabetes and Pregnancy Study Groups (IADPSG) criteria(64). Clinical data were extracted from inpatient medical records, including maternal age, height, pre-pregnancy weight, pre-pregnancy BMI, mode of delivery, gestational age at delivery, neonatal sex, birth weight, LGA or SGA status, NICU admission, HbA1c, and plasma glucose concentrations at 0, 1, and 2 h during the 75-g OGTT at 24–28 weeks of gestation. To minimize confounding effects from pharmacological treatment, all women with GDM received lifestyle intervention only (dietary and exercise management), without any glucose-lowering medications. Participants were excluded if they met any of the following criteria: maternal age ≥35 years; pre-pregnancy BMI ≥28 kg/m²; pre-existing diabetes or a family history of diabetes; prenatal infection; hypertension; kidney disease; thyroid dysfunction; other severe medical conditions during pregnancy; or use of medications that could affect metabolism or inflammation. To specifically investigate the effects of persistent hyperglycemia on placental development and fetal outcomes, GDM participants whose HbA1c decreased to <5.5% after lifestyle management were further excluded(53). After applying these criteria, 52 normal controls and 32 GDM cases were included in the final analysis. Detailed clinical characteristics are provided in FIG.6*A*.

Placental samples were collected within 30 min after delivery. Tissue blocks (∼1 × 1 × 1 cm³) were excised from the maternal side of the placenta near the umbilical cord insertion site, while avoiding areas with calcification, hemorrhage, or necrosis whenever possible. Samples were briefly rinsed with ice-cold phosphate-buffered saline to remove residual blood and amniotic fluid, and non-target tissues were trimmed off. After blotting dry on sterile gauze, each sample was divided into two aliquots (100 mg each). One aliquot was transferred into pre-labeled, pre-chilled cryovials, snap-frozen in liquid nitrogen, and stored at −80°C for protein extraction and downstream analyses. The other aliquot was fixed in 4% paraformaldehyde for 48 h, dehydrated through graded ethanol, cleared, paraffin-embedded, and sectioned serially for immunofluorescence staining.

### Animals

Eight-week-old ICR mice (obtained from Shanghai SLAC Laboratory Animal Co., Shanghai, China; SCXK-2022-0004) were housed in a specific pathogen-free (SPF) facility (12-h light/12-h dark cycle, 24 °C, 50–60% humidity) with ad libitum access to standard chow and water. To obtain timed pregnancies, females were paired with males (2:1 ratio) at 17:00 and checked for vaginal plugs the next morning (∼07:00). The day a plug was observed was designated as gestational day 1 (D1). All animal procedures were approved by the Zhejiang University Institutional Animal Care and Use Committee (Approval number: ZJU20250319) and performed in accordance with institutional guidelines.

### Establishment of the GDM mouse model

Pregnant dams were randomly assigned to vehicle (Veh) or streptozotocin (STZ) groups. On gestational days 6 and 12, dams were fasted for 12 h and then injected intraperitoneally with STZ (100 mg/kg body weight) freshly prepared in 0.1 M citrate buffer (pH 4.5, Solarbio) or with an equal volume of citrate buffer alone (Veh). A 12-hour fasting period was required before each injection. Random blood glucose was measured from the tail vein on D1, D9, and D15 using a handheld glucometer (OneTouch Verio Flex®).

### Glucose monitoring and tolerance test (IPGTT)

IPGTT was performed on D18 (the day of tissue harvest) to assess glucose tolerance. Following an overnight 12-h fast, each dam was injected though intraperitoneal injection with glucose (2 g/kg body weight). Blood glucose was measured from the tail vein at 0, 30, 60, 90, and 120 min after glucose administration use the same glucometer.

### Tissue collection

Pregnant mice were euthanized by cervical dislocation on D18. The uterus was exposed and briefly rinsed in sterile saline. Each uterine horn was opened, and intact conceptuses were gently dissected out. The amniotic membrane was trimmed off the placental surface, and the umbilical cord was cut to separate the placenta and fetus. Placentas and fetuses were immediately weighed, and the fetal-to-placental weight ratio was calculated for each pregnancy. One placenta from each litter was used for subsequent analyses to ensure biological independence. Placental samples for histological analysis were fixed in 4% paraformaldehyde at 4 °C overnight, and samples for molecular assays were snap-frozen in liquid nitrogen and stored at –80 °C until use.

### Litter size recording and birth weight measurement

Pregnant GDM and non-GDM female mice were monitored daily beginning at D18 for delivery. The day of birth was designated as postnatal day 0. Within 15 to 18h after birth, litter size was recorded and each pup was individually weighed using a precision analytical balance. Birth weights were measured in grams (g), and the number of pups per litter was also documented.

### Histological and immunostaining analysis

Fixed placentas were processed and embedded in paraffin using standard procedures. Paraffin blocks were sectioned at 5 μm thickness. For morphological assessment, sections were stained with HE to visualize overall placental structure and zonation, and with PAS reagent to detect glycogen-rich trophoblast cells and any ectopic glycogen deposition. Stained sections were examined under a light microscope (Olympus; BX43). Placental zone organization and boundary integrity were evaluated from H&E images, and PAS staining in the labyrinth was qualitatively compared between groups. Placentas from at least 3 independent pregnancies per group were examined for histological analysis, and representative images are shown.

Paraffin-embedded placental sections were used for IHC and IF staining of specific proteins. For IHC, deparaffinized sections were subjected to heat-induced antigen retrieval in citrate buffer, incubated with primary antibodies at 4°C overnight, and then incubated with an HRP-based secondary antibody followed by DAB chromogen substrate and hematoxylin (Beyotime, C0107) counterstaining. For IF, sections were incubated with primary antibodies at 4°C overnight, followed by fluorophore-conjugated secondary antibodies and nuclear staining with hoechst33342 (Thermo Scientific, H3570). Images were acquired using a Zeiss LSM800 confocal microscope and processed in ImageJ. For each placenta, three sections were analyzed, and three randomly selected fields per section were quantified from anatomically matched regions. Antibodies are listed in SI Appendix, Table S2.

### Lipid droplet staining

The distribution and accumulation of lipid droplets in placental tissues were detected using the lipophilic green fluorescent probe BODIPY 493/503 (Invitrogen, D3922). Fresh placental tissues were embedded in OCT compound, snap-frozen in liquid nitrogen, and sectioned into 8 μm-thick frozen sections using a cryostat. After thawing and drying at room temperature, the sections were fixed with 4% paraformaldehyde for 15 minutes at room temperature. Following three washes with PBS, the sections were incubated with BODIPY 493/503 working solution (1μg/mL) for 30-60 minutes at room temperature in the dark. Subsequently, the sections were washed with PBS and counterstained with Hoechst 33342 solution for 5 minutes at room temperature in the dark to visualize the nuclei. Finally, after washing with PBS and mounting with an anti-fade mounting medium (Solarbio, S2100), the samples were observed and imaged under a laser scanning confocal microscope (Zeiss LSM880). The mean fluorescence intensity of the lipid droplets was quantitatively analyzed using ImageJ software.

### Western blot

Placental tissues were lysed in RIPA buffer (Solarbio, R0020) supplemented with protease and phosphatase inhibitor cocktails (Solarbio, IKM1020) and PMSF (Solarbio, P0100). After centrifugation at 4°C, protein concentration was quantified using the Pierce BCA Protein Assay Kit (Thermo Scientific, 23225). Samples were denatured in 5× loading buffer containing DTT (Solarbio, P1040) at 100°C for 10 min, separated by SDS-PAGE, and transferred to polyvinylidene fluoride (PVDF) membrane (IPVH00010, Merck). PVDF membranes were blocked in 5% skim milk in TBST for 1 h at room temperature and incubated with primary antibodies at 4°C overnight. After washing, membranes were incubated with HRP-conjugated secondary antibodies for 1 h at room temperature, and signals were detected using enhanced chemiluminescence (Millipore, WBKLS0500) and visualized with a FUSION FX6 imaging system equipped with Image Lab Software (Vilber). Band intensities were quantified in ImageJ and normalized to H3 or β-actin. Antibodies are provided in SI Appendix, Table S1.

### Metabolomic analysis of placental tissues

Placental tissues were collected at D18. The tissues were immediately weighed and placed in 80% LC-MS-grade methanol at a ratio of 100 mg/mL. Placental tissues were homogenized using a Cryogenic Tissue Grinder (Jingxin, JXFSTPRP-CL-BSC) at 4 °C. The homogenized tissue–methanol mixtures were flash-frozen in liquid nitrogen and stored at −80 °C until further processing. Samples were transported to Shanghai Applied Protein Technology on dry ice.

At the company facility, frozen tissue–methanol mixtures were sonicated at 4 °C, followed by incubation (SCIENTZ, SCIENTZ08-IIIC) at −20 °C for 10 min. Samples were then centrifuged at 14,000 × g for 20 min, and the supernatants were collected, dried, and reconstituted in 50% acetonitrile (ACN) for LC–MS analysis. Samples were analyzed in randomized order, and pooled quality-control (QC) samples were injected throughout the analytical run to monitor system stability. LC–MS/MS raw data were processed using the company’s standard workflows for peak detection and alignment, and metabolite intensities were normalized prior to statistical analysis. The resulting metabolite abundance matrix was returned for downstream differential analysis. Differential metabolite analysis was performed using Student’s t-test. Metabolites with a fold change >1.2 and *P* < 0.05 (two-sided) were considered significantly altered.

### Enzyme-linked immunosorbent assay (ELISA) for acetyl-CoA

Placental acetyl-CoA levels were measured using a commercial acetyl-CoA ELISA kit (Sangon Biotech, D751001). Frozen placental tissue was homogenized in the kit extraction buffer, and the assay was carried out according to the manufacturer’s instructions. In brief, standards and diluted samples were added to a 96-well microplate pre-coated with capture antibody, and through a competitive ELISA mechanism the acetyl-CoA concentration in each sample was determined. After the final color development step, absorbance at 450 nm was measured on an absorbance microplate reader (TECAN, Sunrise™). A standard curve was used to calculate acetyl-CoA concentrations in pmol per mg of tissue. Each sample was measured in duplicate, and technical replicates were averaged.

### NAD⁺/NADH and NADP⁺/NADPH assays

The levels of NAD and NADP in placental tissue were quantified using colorimetric assay kits (Beyotime, S0175, S0179). Placental samples (10–20 mg of tissue per sample) from each dam were processed according to the kit protocols. Briefly, tissues were lysed in the provided extraction buffer and the extracts were filtered or clarified. For each sample, two parallel reactions were set up to measure the total amount of NAD (NAD^+^ + NADH) and the amount of NADH alone; the difference represents NAD^+^. Similarly, total NADP and NADPH were measured in parallel. The assay utilizes an enzyme cycling reaction that produces a colorimetric product detectable at 450 nm proportional to the NAD(H)/NADP(H) content. Absorbance was read at 450 nm on a microplate reader, and concentrations were calculated from standard curves. Values were normalized to the tissue input (results expressed as pmol /mg tissue).

### RNA isolation, reverse transcription, and quantitative real-time PCR (qPCR)

Total RNA was extracted using TRIzol (Thermo Fisher, 15596018CN). The OD260/280 ratio was measured using a NanoDrop 2000 spectrophotometer (Thermo Fisher Scientific) to evaluate RNA purity. cDNA was synthesized from 1 μg RNA using a reverse transcription kit (Vazyme, R223-01). Quantitative PCR was performed with SYBR master mix (Vazyme, Q711) on a Real-Time PCR Detection System (Thermo Fisher Scientific, QuantStudio™ 3). Relative expression was calculated using the 2^−ΔΔCt method with Actb as internal controls. Primer sequences are provided in SI Appendix, Table S2.

### Bulk RNA-seq and data analysis

Bulk RNA-seq was performed using total RNA extracted from whole placentas collected at D18. Total RNA was isolated as described above and transported to Shanghai Applied Protein Technology on dry ice for library preparation and sequencing. Specifically, Poly(A)+ mRNA was enriched using oligo(dT) magnetic beads. The paired-end libraries were generated using the VAHTS Universal V6 RNA-seq Library Prep Kit. Libraries were sequenced on the DNBSEQ-T7 platform (Shanghai Applied Protein Technology) to produce 150-bp paired-end reads.

RNA-seq data were analyzed as previously described. Briefly, raw sequencing reads were processed using Trim Galore (https://github.com/FelixKrueger/TrimGalore) to remove adapter sequences and low-quality reads. Clean reads were aligned to the mouse reference genome (GRCm39/mm39) using HISAT2 (version 2.2.1). The FPKM (Fragments Per Kilobase of transcript sequence per Million base pairs sequenced) values of each gene in every sample were calculated using the FeatureCounts software. Differential expression analysis between the Veh and STZ groups was performed using the edgeR. Genes with a fold change greater than 1.5 and *P* < 0.05 were considered DEGs. GO enrichment analysis was conducted using DAVID (https://david.ncifcrf.gov), with corrected *P* < 0.05 considered statistically significant. GSEA was performed using the GSEA software (http://www.broadinstitute.org/gsea/index.jsp). In addition to the gene set related to trophoblast cell differentiation as defined by the literature(65), all the remaining gene sets were obtained from the Molecular Signatures Database (MSigDB).

### Statistics and reproducibility

Statistical analyses were performed using GraphPad Prism (v 10.6.0). Key experiments were independently repeated at least three times. For two-group comparisons, two-tailed unpaired Student’s t tests were used unless otherwise noted. For multiple groups, one-way ANOVA with Tukey’s multiple-comparison test was applied. For time-course glucose measurements, two-way ANOVA with appropriate multiple-comparison correction was used. Data are presented as mean ± SEM (or mean, consistent with figure legends). *P* < 0.05 was considered statistically significant.

---

**Fig. S1.**
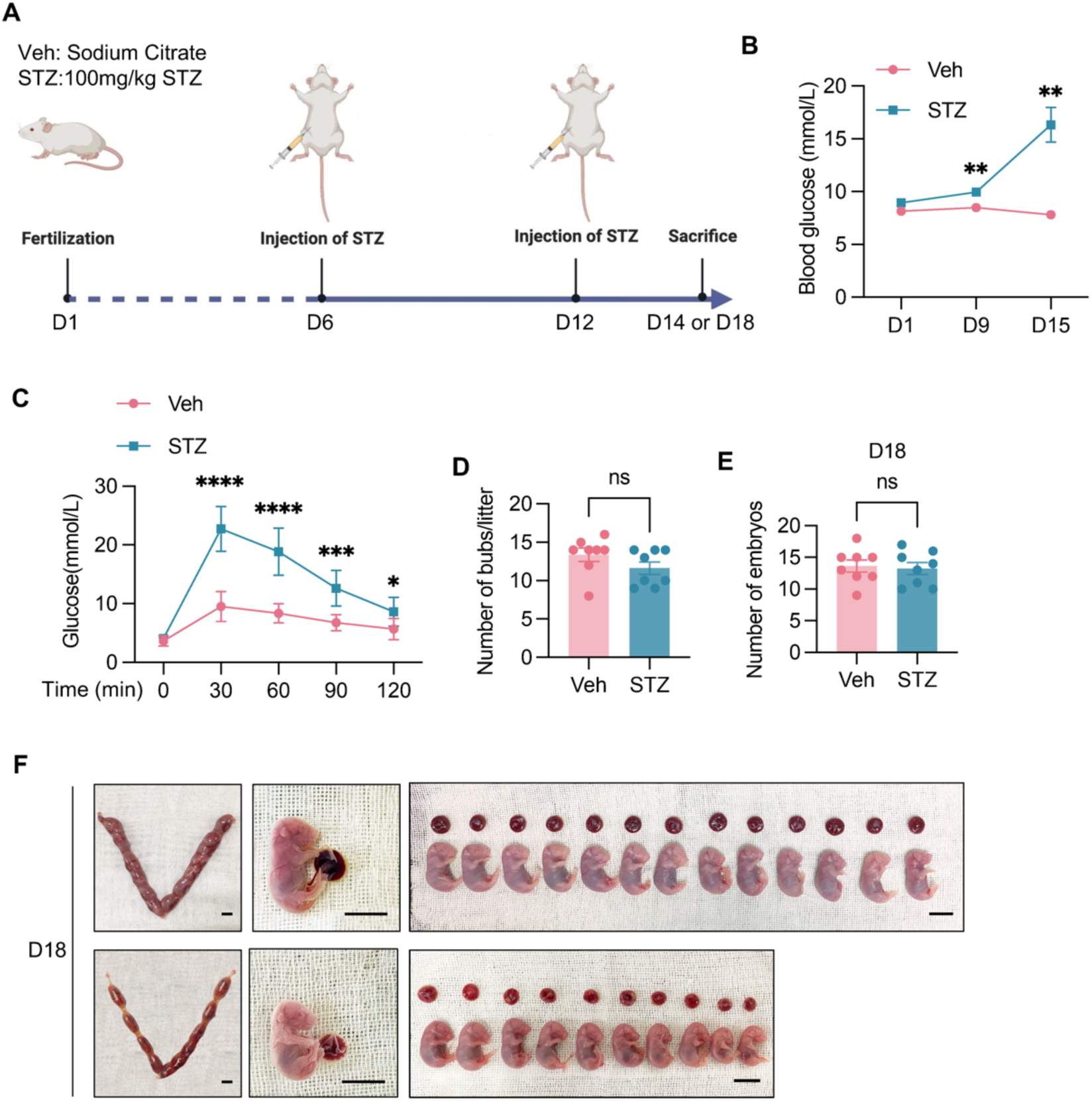
Successful establishment of a mid- to late-gestation hyperglycemic GDM mouse model. (***A***) Experimental schematic for STZ-induced gestational hyperglycemia in pregnant mice. (***B***) Blood glucose levels measured at indicated gestational time points in Veh and STZ. Data represent the mean ± SEM. Two-way ANOVA. \*\**P*<0.01. (***C***) Glucose tolerance test (GTT) in Veh and STZ pregnancies. Two-way ANOVA. Data are mean ± SEM. \**P* < 0.05, \*\*\**P* < 0.001, \*\*\*\**P* < 0.0001. (***D, E***) Litter size (number of pups/litter) (*D*) and number of embryos on D18 (*E*) in Veh and STZ pregnancies. Two-tailed unpaired Student’s t-test, ns, not significant. \*\*\*\**P* < 0.0001. **(*F*)** Representative images of gravid uteri and embryos from Veh and STZ pregnancies on D18. Scale bar, 1cm.

**Fig. S2.**
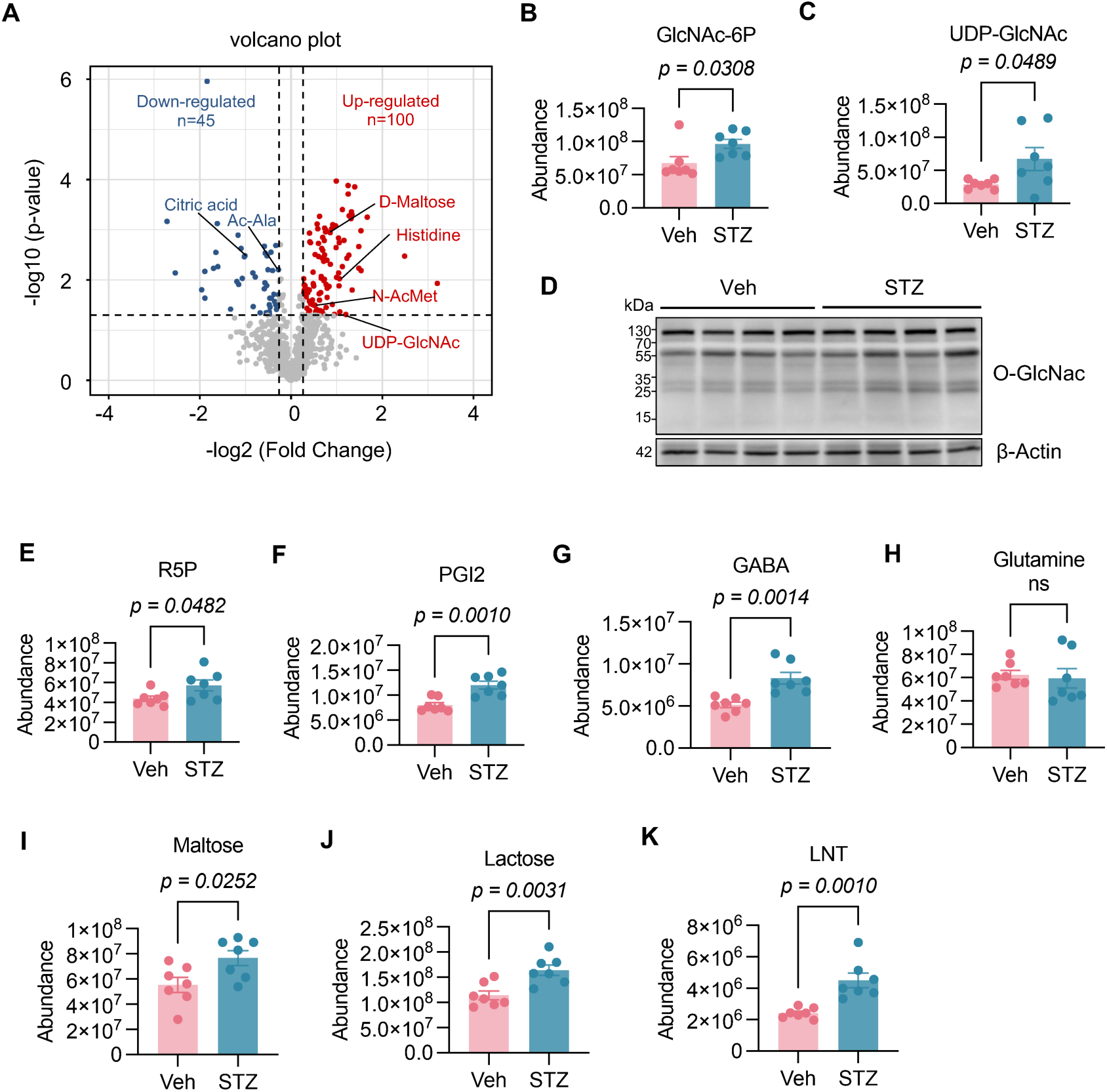
Untargeted metabolomics identifies activation of hexosamine and carbohydrate biosynthesis pathways in GDM placenta. (***A***) Volcano plot of differential metabolites in Veh and STZ placentas on D18. (***B, C***) Quantification of GlcNAc-6P (*B*) and UDP-GlcNAc (*C*) in Veh and STZ placentas on D18 detected by LC-MS. (***D***) Immunoblotting analysis of global protein O-GlcNAc in Veh and STZ placentas with β-actin as a loading control. (***E–K***) Quantification of representative metabolites in Veh and STZ placentas on D18 detected by LC-MS. Data represent the mean ± SEM. Two-tailed unpaired Student’s t-test.

**Fig. S3.**
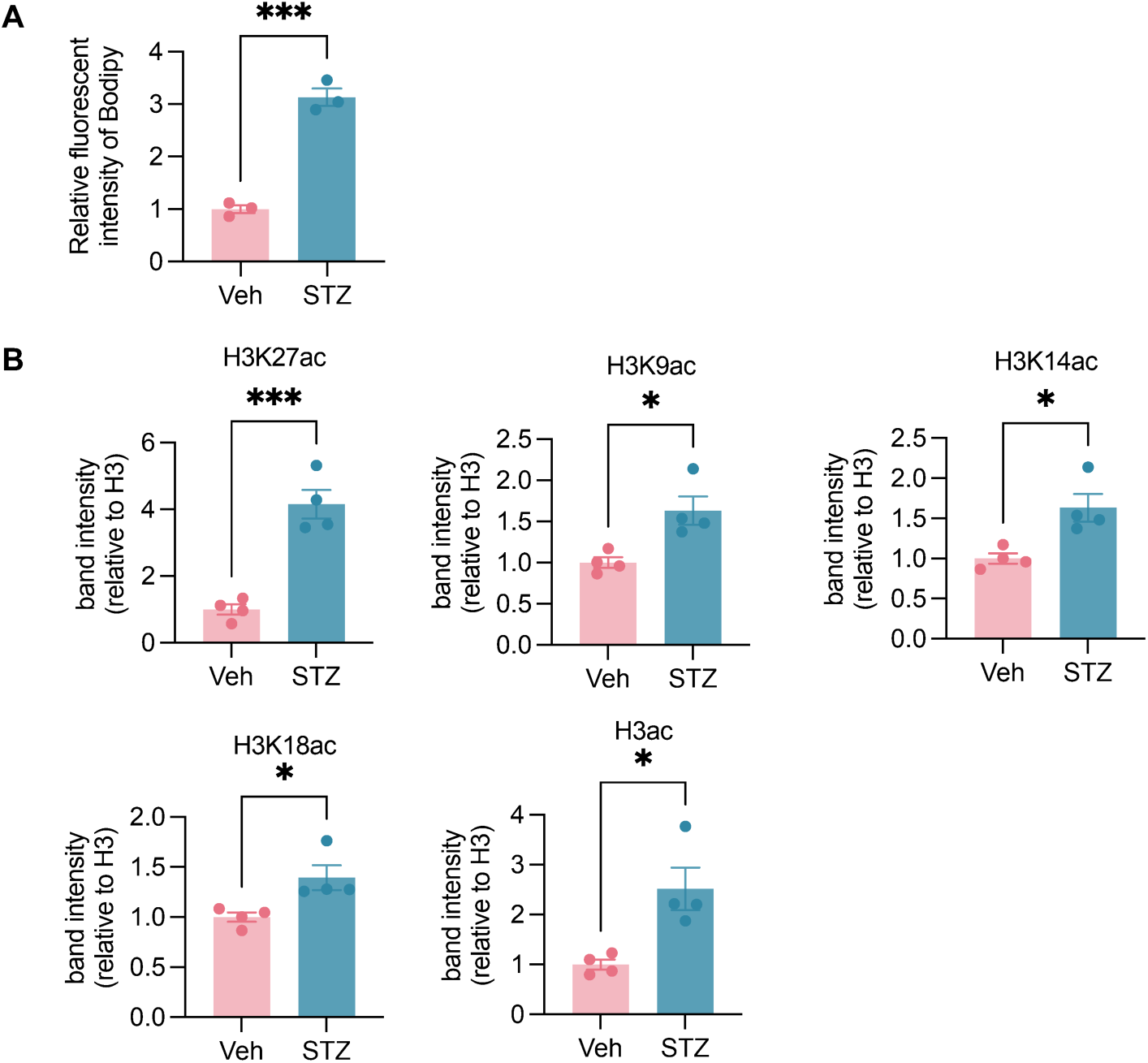
Quantification of lipid accumulation and histone acetylation changes in GDM placenta. (*A*) Quantification of relative fluorescent intensity of BODIPY staining in Veh and STZ placentas on D18. **(*B*)** Densitometric quantification of histone acetylation marks normalized to total H3 in Veh and STZ placentas on D18. Data represent the mean ± SEM. Two-tailed unpaired Student’s t-test, \**P* < 0.05, \*\*\**P* < 0.001.

**Fig. S4.**
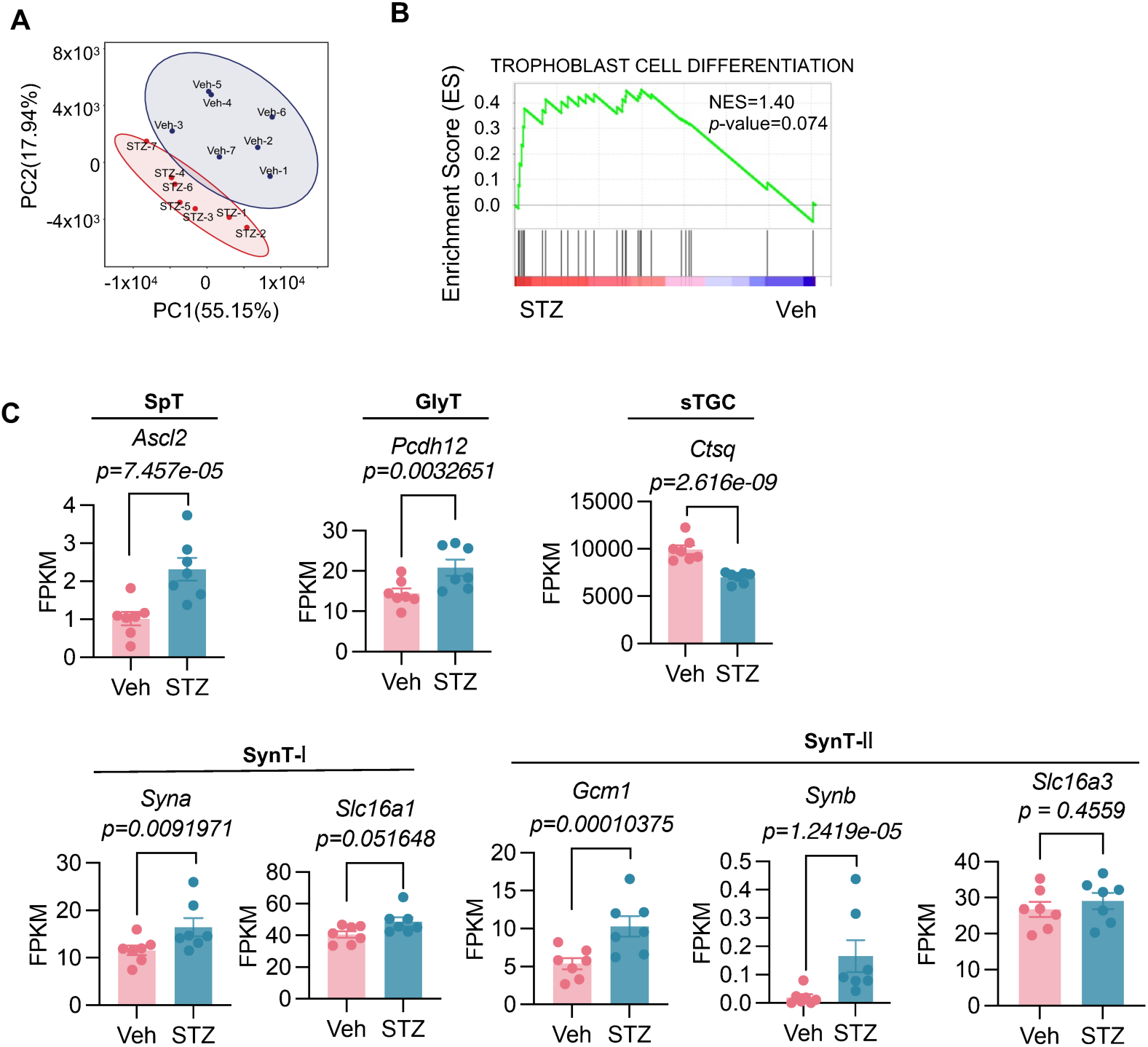
Transcriptomic analyses support enhanced trophoblast differentiation programs in GDM placenta. (*A*) PCA of transcriptomes from Veh and STZ placentas on D18. **(*B*)** GSEA analysis on the gene set related to trophoblast differentiation of Veh and STZ placentas on D18. **(*C*)** FPKM (Fragments Per Kilobase per Million) of representative trophoblast lineage markers (SpT, GlyT, sTGC, SynT-I, SynT-II) in Veh and STZ placentas on D18.

**Fig. S5.**
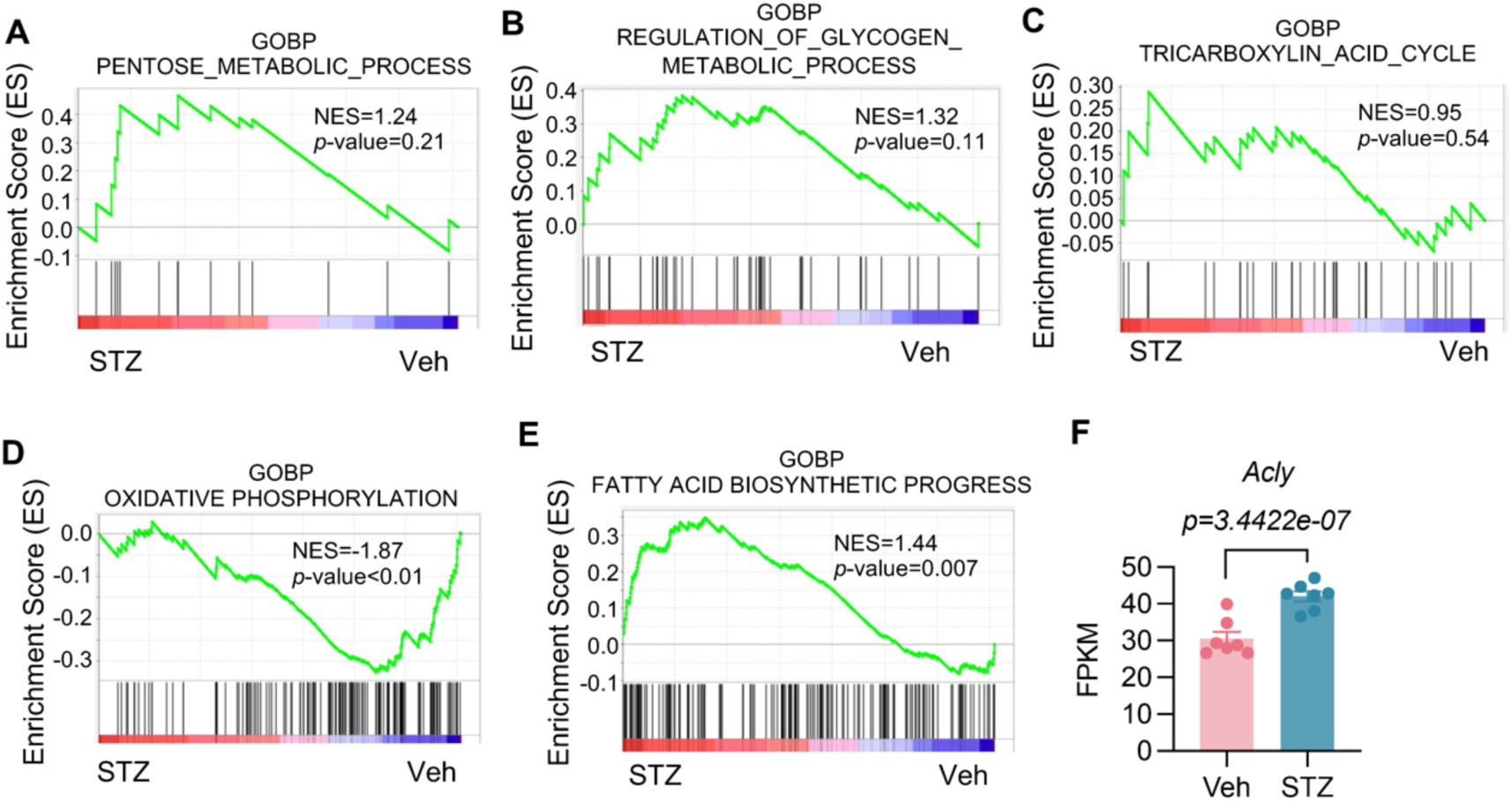
GSEA indicates activation of fatty acid biosynthesis and suppression of OXPHOS in STZ placentas. (*A*–*E*) GSEA analysis on specific metabolic pathway of Veh and STZ placentas on D18. **(*F*)** RNA-seq quantification of *Acly* expression (FPKM) in Veh and STZ placentas on D18.

**Fig. S6.**
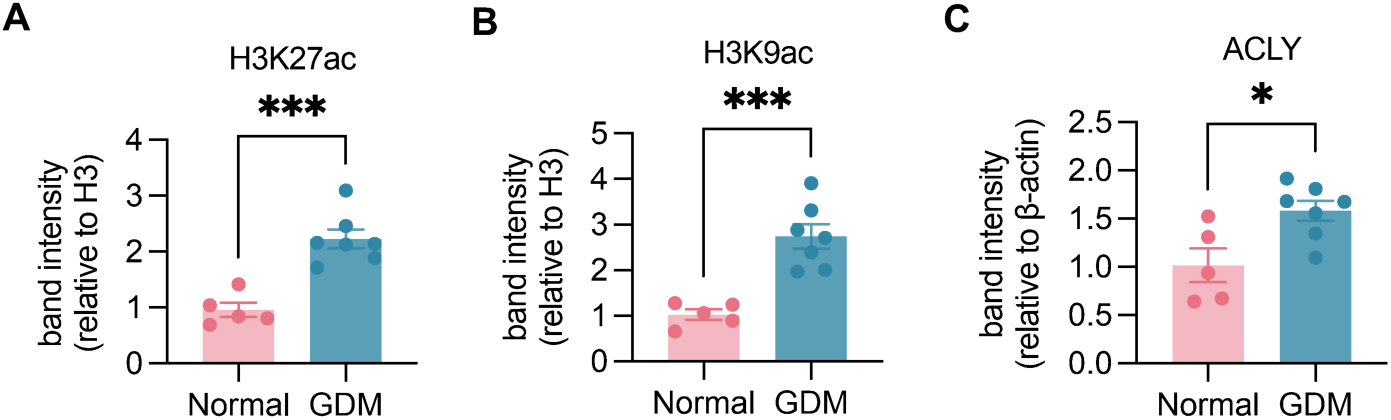
Quantification of histone acetylation and ACLY changes in GDM placenta. (*A-C*) Densitometric quantification of H3K27ac (*A*) and H3K9ac (*B*) normalized to total H3 and ACLY (*C*) normalized to β-actin. Data represent the mean ± SEM. Two-tailed unpaired Student’s t-test, \**P* < 0.05, \*\*\**P* < 0.001.

