## Supplementary material for "Metabolic adaptation to maternal hyperglycemia via ACLY-dependent acetyl-CoA production drives epigenetic remodeling and dysregulated placental development": Table S1

**Table S1.** Antibodies used in this study.

| Antibody | Catalog number | Final concentration | Manufacturer | Host species |
| --- | --- | --- | --- | --- |
| β-actin | 66009-1-Ig | 1:20000 (WB) | Proteintech | Mouse |
| ACLY | ab40793 | 1:10000 (WB)<br>1:50 (IF) | Abcam | Rabbit |
| H3K9AC | 9649T | 1:1000 (WB) | Cell Signaling Technology | Rabbit |
| H3K14AC | 7627S | 1:1000 (WB) | Cell Signaling Technology | Rabbit |
| H3K18AC | 13998T | 1:1000 (WB) | Cell Signaling Technology | Rabbit |
| H3K27AC | 8173S | 1:1000 (WB)<br>1:200 (IF) | Cell Signaling Technology | Rabbit |
| H3AC | 61638 | 1:5000 (WB) | Active Motif | Rabbit |
| Histone H3 | 4499T | 1:2000 (WB) | Cell Signaling Technology | Rabbit |
| O-Linked N-Acetylglucosamine | ab2739 | 1:1000 (WB) | Cell Signaling Technology | Mouse |
| CD31 | ab281583 | 1:4000 (IHC) | Abcam | Rabbit |
| MCT1 | AB1286-I | 1:400 (IF) | Sigma | |
| MCT4 | sc-376140 | 1:400 (IF) | Santa Cruz Biotechnology | |
| hCG | ab9582 | 1:200 (IF) | Abcam | |
