## Supplementary material for "Metabolic adaptation to maternal hyperglycemia via ACLY-dependent acetyl-CoA production drives epigenetic remodeling and dysregulated placental development": Table S2

**Table S2.** Sequence of the primers for qPCR used in this study.

| Mouse | Forward Primer (5'-3') | Reverse Primer (5'-3') |
|---|---|---|
| *Acly* | AGGAAGTGCCACCTCCAACAGT | CGCTCATCACAGATGCTGGTCA |
| *Ascl2* | TTTCCTGTGCCGCACCAGAACT | CAGCGACTCCAGACGAGGTGG |
| *Ctsq* | TCTGGAGGCTCAGGCAACCTAT | GTTGCTACAGCATCCATGAGGAC |
| *Gcm1* | AGCAGAGGCAAGAAGAGCCATG | TTGGACGCCTTCCTGGAATGAC |
| *Syna* | CTTTCCAAGGCTCTCTCGGACA | CTCAGCCACAATGAGGTCCAGA |
| *Synb* | CAAACACTGCCATACCTCTCCG | CACTGACATGGTAACAGGGTGG |
| *Slc16a1* | GACCATTGTGGAATGCTGCCCT | CGATGATGAGGATCACGCCACA |
| *Slc16a3* | TCCATCCTGCTGGCTATGCTCT | CAGAAGGACGCAGCCACCATTC |
| *Prl2c2* | TTCCTTCCAACTCCAGAAAACAAG | CTAGATCGTCCAGAGGGCTTTC |
| *Pcdh12* | TTCTGAGGAGCCTGGTTAGGCT | CAAGGACAGCAGCTGGGAGATT |
| *β-actin* | AGTGTGACGTTGACATCCGT | GCAGCTCAGTAACAGTCCGC |
